# Multiplex *in situ* hybridization within a single transcript: RNAscope reveals dystrophin mRNA dynamics

**DOI:** 10.1101/791780

**Authors:** J.C.W. Hildyard, F. Rawson, D.J. Wells, R.J. Piercy

## Abstract

Dystrophin plays a vital role in maintaining muscle health, yet low mRNA expression, lengthy transcription time and the limitations of traditional *in-situ* hybridization (ISH) methodologies mean that the dynamics of dystrophin transcription remain poorly understood. RNAscope is highly sensitive ISH method that can be multiplexed, allowing detection of individual transcripts at sub-cellular resolution, with different target mRNAs assigned to distinct fluorophores. We present a novel approach, instead using RNAscope probes targeted to 5’ and 3’ regions of the same transcript: labelling muscle dystrophin mRNA in this manner allows transcriptional dynamics to be deciphered in health and disease, resolving both nascent myonuclear transcripts and exported mature mRNAs (the latter absent in dystrophic muscle, yet restored following therapeutic intervention). We show that even in healthy muscle, immature dystrophin mRNA predominates (60-80% of total), with the surprising implication that the half-life of a mature transcript is markedly shorter than the time invested in transcription: at the transcript level, supply may exceed demand. Our findings provide unique spatiotemporal insight into the behaviour of this long transcript (with implications for therapeutic approaches), and further suggests this modified multiplex ISH approach is well-suited to long genes, offering a highly tractable means to reveal complex transcriptional dynamics.

## Introduction

The dystrophin gene is the largest in the genome: at approximately 2.4Mbp in length this single gene accounts for almost 0.1% of human haploid DNA and occupies fully 1.5% of the X chromosome where it resides. Transcription of a single full-length 79-exon dystrophin mRNA is estimated to take 16 hours, with translation giving rise to dp427 dystrophin (a protein product of 427kDa). This lengthy gene also displays unconventional transcriptional behaviour: the dp427 transcript can arise from three separate promoters controlling cortex, muscle and Purkinje cell expression, producing dp427c, dp427m and dp427p respectively (1, 2). Each promoter confers a unique first exon but the other 78 are shared, resulting in dp427 proteins essentially identical in sequence, differing only in the first 3-11 amino acids (of a total of almost 3700). The dystrophin gene also carries several internal promoters producing shorter transcripts, generating multiple N-terminally truncated protein isoforms (similarly named by molecular weight, thus dp260 (3), dp140 (4), dp116 (5), and dp71 (6)): see Figure 1A. These isoforms show clear tissue-specific expression patterns and likely play unique cellular roles, but most studies to date have focussed on dp427m, the full-length muscle isoform: mutations almost anywhere in the dystrophin gene affect this isoform, and this isoform is critical to human health.

**Figure 1:**
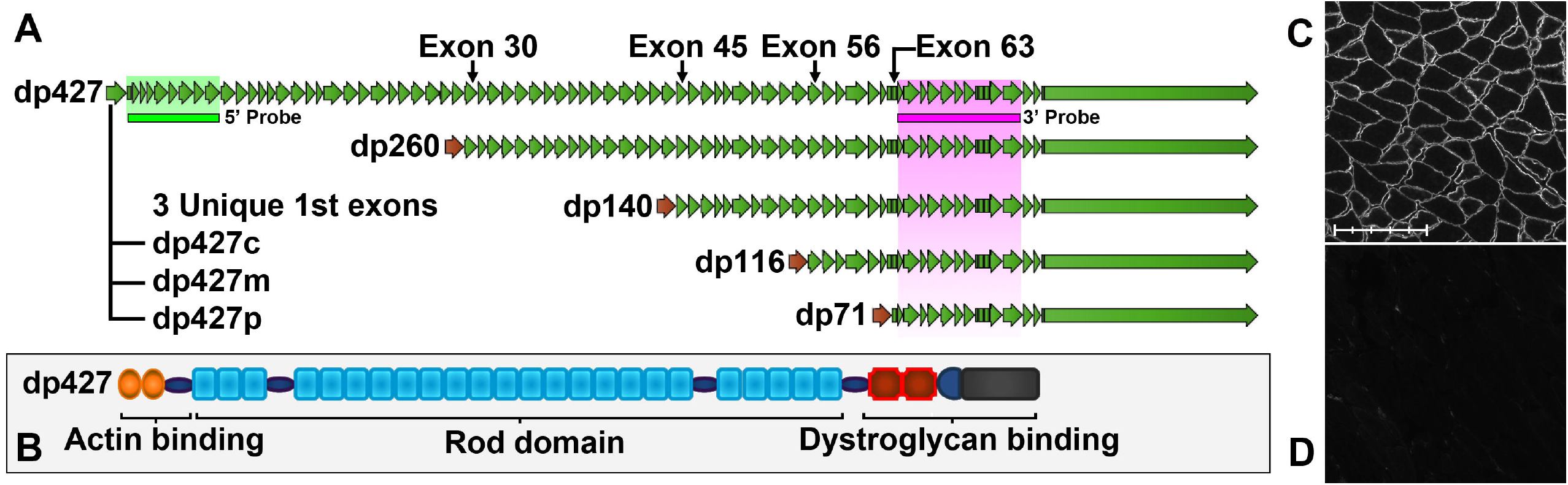
Dystrophin mRNA isoforms and full-length protein. (A) Cortical, muscle and Purkinje isoforms of full length dystrophin (dp427) have unique first exons but share all 78 downstream exons. Dp260 shares all sequence from exon 30, dp140 from exon 45, dp116 from exon 56 and dp71 from exon 63. Target sequence regions of the 5’ and 3’ probes used in this study are shown (green and magenta bars): 5’ probe recognizes dp427 only (c, m and p), while 3’ probe will recognize all dystrophin isoforms. (B) dp427 protein has three principal domains: N-terminal actin binding, spectrin repeat rod domain, and C-terminal dystroglycan binding. Skeletal muscle dystrophin protein localizes to the sarcolemma in healthy mouse quadriceps muscle (C) but is absent in dystrophic mdx quadriceps muscle (D). Scale bar: 200μm (subdivisions: 40μm).

Dp427m is the only dystrophin isoform expressed in skeletal muscle (7), and the long dp427 protein it encodes contains three principal domains: an N-terminal actin-binding region, a C-terminal dystroglycan-binding region, and a lengthy ‘rod’ domain of 24 spectrin-like repeats linking the two (Figure 1B) (8). This protein localizes to the sarcolemma (Figure 1C) where it forms a key component of the dystrophin-associated glycoprotein complex (DAGC), localizing proteins that aid muscle function such as neuronal nitric oxide synthase (nNOS) (9) and acting as a physical link between the intracellular actin cytoskeleton and the extracellular matrix environment (10). Insufficient or absent muscle dp427 (Figure 1D) results in weakness of the muscle sarcolemma, leaving muscle fibres vulnerable to contraction-induced injury, leading to the incurable and fatal muscle-wasting disease, Duchenne muscular dystrophy (DMD). This X-linked disease, affecting 1 in 5000 new born boys (11, 12), is characterized by cycles of muscle degeneration and compensatory regeneration, with concomitant persistent inflammation and progressive loss of muscle tissue to fat and fibrotic scarring: boys become wheelchair-bound between the age of 8 and 12, and death typically occurs in the 20s-30s via either cardiac or respiratory failure (13). Mutations that cause premature termination of dp427 result in DMD, however mutations that cause in-frame internal truncation of dp427 typically lead to the milder condition, Becker muscular dystrophy (BMD), suggesting that while N and C termini are critical for function, the rod domain is largely dispensable: indeed, use of antisense oligonucleotides to mask splicing sites in the dp427 pre-mRNA leads to omission of targeted exons in DMD-causing transcripts and consequent restoration of reading frame, offering a means to change DMD to a BMD phenotype. These ‘exon-skipping’ approaches represent a promising therapeutic approach.

Dystrophin remains a challenging target of investigation: dp427 represents only a tiny fraction of total muscle protein (~0.002% (14)), and levels of transcripts within skeletal muscle are consequently similarly modest (15, 16). Immunohistochemistry with specific antibodies allows the subcellular localization of dystrophin protein to be determined with high accuracy, but similar studies at the mRNA level are more challenging. *In-situ* hybridization (ISH) with biotinylated (17) or radiolabelled (18) probes suggests a sarcolemmal localization of the dp427 transcript, but further details are confounded by the innate limitations of this technique and the low transcript abundance. Most studies of dp427 mRNA dynamics have therefore typically measured dystrophin expression in whole muscle extracts, necessarily discarding spatial information. As muscle tissue is host to multiple cell types (19), such approaches also introduce unavoidable noise (a problem exacerbated in dystrophic muscle where infiltrating inflammatory cells, fibroblasts, adipocytes and differentiating myoblasts further diversify the cellular milieu (20)). Despite these limitations, pioneering work by Tennyson *et al* (21) determined estimates for dp427 transcription time of the order of 16 hours, and showed that splicing occurred co-transcriptionally (work subsequently supported by others investigating transcription of long genes (22) where co-transcriptional splicing and elongation rates of 40-60 bases per second were reported). Further investigations (16) suggested that in human muscle dp427 5’ sequence was present in excess of 3’ sequence (i.e. nascent transcripts exceeded mature), suggesting the half-life of mature dp427 might be modest, but at the time such work necessitated transcript estimation via somewhat crude RT-PCR autoradiography. An elegant recent study by Gazzoli *et al* revealed that co-transcriptional splicing of dp427 occurs non-sequentially, and that long introns in particular (several are >100kb) are spliced in multiple steps (23), however most studies of dp427 focus on disease-or therapeutic-relevant metrics such as the efficiency of exon-skipping (percentage of skipped transcripts) rather than more general properties of the mRNA itself (particularly at the histological level). Recent technological advances such as RNAscope (24) have led to a resurgence in ISH-based investigations, suggesting that detailed study of muscle dp427 mRNA at the histological level might now not only be practical, but also (given the large size and low abundance of the transcript, and the unique multinucleate nature of myofibres) highly nuanced.

The RNAscope ISH approach employs proprietary ‘ZZ’ oligonucleotide probe pairs: each probe carries 18-25 bases of target-complementary sequence linked to one half of a preamplifier-binding motif. The preamplifier carries multiple amplifier-binding motifs, and the amplifiers in turn are able to bind peroxidase enzymes in very high local concentrations. Use of 20 such ZZ pairs in series allows targeting of ~1000 bases of target sequence (rendering this method highly specific), and use of the amplification strategy with peroxidase-activated tyramide dyes affords very high sensitivity, allowing detection of individual mRNA transcripts with sub-cellular resolution (Figure 2A-E). Crucially, this technique can also be multiplexed: by use of probe-specific amplifiers and sequential labelling, multiple mRNA species can be resolved, each with a different fluorophore (Figure 2F). Our approach is a further refinement of this method: we reasoned that the substantial size of the full-length dystrophin transcript would readily permit multiple 20ZZ probe sets to bind to the same mRNA, a hypothesis supported by Smith *et al*, using conventional fluorescent probes (25) to resolve the nascent transcriptional locus within cultured myonuclei. Using probes targeted to 5’ and 3’ sequence of dp427 (see Figure 1A) offers an elegant means to label transcripts in a temporal fashion (in a manner similar to that first used by Garcia *et al* (26), but without recourse to complex transgenics), and should permit unprecedented resolution of individual mature and nascent dystrophin mRNA molecules within skeletal muscle: nascent transcripts would bind 5’ only, while mature would bind both (Figure 2G). Here, we have used these probes in healthy and dystrophic (*mdx*) mouse muscle and in *mdx* muscle rescued with exon-skipping agents to reveal remarkable insights into the localization and dynamics of this critical transcript in health and disease. Our work supports the findings of previous investigations while adding valuable spatiotemporal refinements unachievable by other means, and suggests that this approach offers a powerful and versatile new tool not only for the study of dystrophin, but for other transcripts produced from long genes.

**Figure 2:**
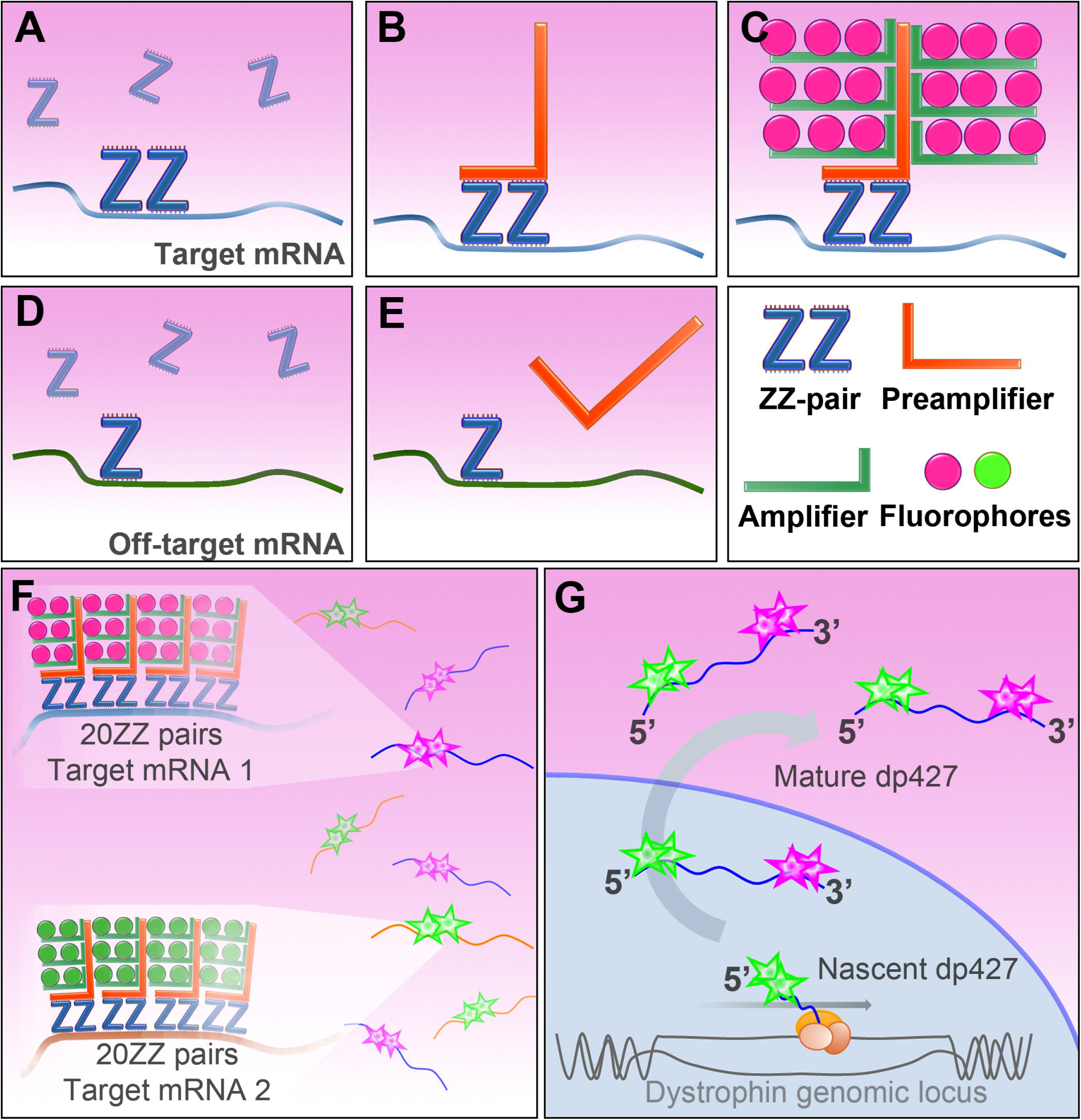
RNAscope ISH. (A) RNAscope ZZ probe pairs bind 30-50 bases of target sequence. Adjacent probes create a target motif for preamplifier molecules (B), which bind amplifiers, which bind probe-specific enzyme to deposit high local concentrations of fluorophore (C). Off-target effects are minimized as both elements of the ZZ pair must bind in close proximity. A single Z probe is insufficient to bind preamplifier (E). Probe sets of 20 sequential ZZ pairs (F) can be used to ensure high specificity, and sequential probe-specific dye-labelling allows different fluorophores to label different mRNA target molecules. (G) dp427 can be targeted with multiple probes: sets targeting 5’ (exons 2-10) and 3’ (exons 64-75) regions of dp427 permit distinction of nascent (5’ only) and mature (5’ and 3’) transcripts.

## Methods

### Probe design

20ZZ RNAscope probes (ACDBio) were designed to mouse dystrophin sequence (accession number NM_007868.6). The catalogue probe (Mm-Dmd, Cat. No. 452801) recognizes residues 320-1295 (exons 2 to 10) of the full-length (dp427) dystrophin transcript. This probe (henceforth: *5’probe)* recognizes full-length dp427 only, but will bind both nascent and mature dp427 transcripts. A further custom probe (Mm-Dmd-O1, henceforth: *3’probe*) was designed to residues 9581-10846 (exons 64-75). This 3’ probe hybridizes to sequence emerging late in transcription, favouring mature transcripts. Given shared sequence, 3’ probe will recognize essentially all dystrophin isoforms (dp427, 260, 140, 116, 71 –see figure 1A).

Positive control probes to POLR2A (NM_009089.2, residues 2802-3678), PPIB (NM_011149.2, residues 98-856) and UBC (NM_019639.4, residues 36-860) were used to confirm preservation of sample RNA, while negative control probes to bacterial DapB (EF191515, residues 414-862) were used to establish nonspecific labelling.

### Sample preparation

#### Animals

Seven 40-week old male mice (3 *mdx* and 4 wild-type C57BL/6J −referred to within the text as *mdx* 1-3 and WT 1-4) bred under UK Home Office Project Licence PPL 70/7777 were killed by cervical dislocation, with tissues harvested rapidly post mortem. Quadriceps muscles from a further three 32-week old male *mdx* mice (untreated *–mdx* 4 within the text-, or treated intravenously with PIP6a peptide-conjugated antisense morpholino oligonucleotide (PPMO) at 6mg.kg^−1^ and 12.5mg.kg^−1^) were archive samples taken from two separate studies ((27) and manuscript in preparation). Briefly, mice were treated via tail vein injection with the PPMO starting at 12 weeks of age, continuing for a total of 10 injections each separated by two weeks with collection two weeks after the last dose. Quadriceps muscles were mounted longitudinally (a configuration that generates longitudinal and transverse regions within a single section) in cryoMbed mounting medium (Bright instruments Ltd) and flash-frozen in liquid N_2_-cooled isopentane. Freezing in an essentially relaxed configuration rather than pinned out to L0 reduces fibre hypercontraction immediately post-sectioning (see below).

#### Cryosectioning

muscle tissues were sectioned at −25°C to 8μm thickness using an OTF5000 cryostat (Bright) and mounted on glass slides (SuperFrost, VWR). Preparation of longitudinal sections from frozen mouse muscle is non-trivial: transient melting during transfer of section to slide typically results in considerable fibre hypercontraction as contractile proteins respond to abundant free calcium ions.

RNAscope ISH involves multiple detergent washes and incubations at 40°C, necessitating the enhanced tissue affinity of SuperFrost slides, but this slide format is incompatible with EDTA-coating methods of Pearson and Sabarra (28). It was determined that freezing muscles in a relaxed state and sectioning at 8 micron thickness generates acceptable longitudinal sections without the requirement for EDTA: at this thickness, shear forces generated by contractile proteins in the presence of free Ca^2+^ appear largely insufficient to overcome the rapid adhesion to SuperFrost slides. Serial sections were collected and slides were air-dried at −20°C for 1 hour before storage at −80°C until use.

### Staining

#### Immunofluorescence

slides were allowed to equilibrate to room temperature, blocked for 1hr using 5% milk powder (Marvel) in PBS + 0.05% tween (PBS-T), and then co-immunolabelled with rabbit polyclonal anti-dystrophin (ab15277, Abcam, 1:800) and rat monoclonal anti-perlecan (clone A7L6, Thermofisher, 1:2000), in PBS-T for 1hr. Secondary labelling used Alexa-fluor conjugated antibodies (anti-rabbit 488 and anti-rat 594 respectively, Thermofisher, both 1:1000, 1hr). Nuclei were stained using Hoechst 33342 (Thermofisher, 1:2000, 10 minutes), and slides were then washed and mounted using Hydromount (National Diagnostics).

#### SDH

slides were allowed to equilibrate as above, then incubated with freshly-prepared SDH staining solution (125mM sodium succinate, 1.5mM nitroblue tetrazolium, 60mM Tris HCL, pH 7.0) at 37°C for 1 hour, fixed in formal calcium for 15mins at room temperature, then washed and mounted in Hydromount.

### RNAscope slide preparation

The combination of protease digestion and detergent-rich wash buffer used by RNAscope is incompatible with fresh-frozen muscle prepared according to standard protocols: while connective tissue/extracellular matrix elements (and most nuclei) were readily retained throughout the protocol, myofibrillar content was rapidly lost leaving empty myofibre ‘ghosts’ (see Supplementary Figure S1). Preparation of frozen muscle thus required extended fixation and an additional baking step (both indicated with an asterisk): slides were removed from −80°C storage and placed immediately into cold (4°C) 10% neutral-buffered formalin, then incubated at 4°C for 1 hour*. Slides were then dehydrated in graded ethanols (50%, 70%, 100% x2, 5mins in each, room temperature) and left in 100% ethanol at −20°C overnight, then air-dried and baked at 37°C for 1 hour*. Sections were ringed using hydrophobic barrier pen (Immedge, Vector Labs) and then treated with RNAscope hydrogen peroxide (ACDbio) for 15mins at room temperature to quench endogenous peroxidase activity. After washing twice in PBS, slides were protease-treated (RNAscope Protease IV, ACDbio) for 30mins at room temperature and washed a further two times in PBS before use in RNAscope multiplex assay (see below).

### RNAscope multiplex assay

Multiplex assays were performed according to the manufacturer’s protocols. All incubations were at 40°C and used a humidity control chamber (HybEZ oven, ACDbio). Probe mixes used were as follows:

- RNAscope 3-plex positive control probe set (320881): POLR2A, PPIB and UBC (low, moderate and high expression targets supplied as C1, C2 and C3 probes, respectively)
- RNAscope 3-plex negative control probe set (320871): Bacterial DapB (in C1, C2 and C3 probe sets)
- RNAscope mouse dystrophin probe set: Mm-Dmd (452801) and Mm-Dmd-O1 (custom probe) (C1 and C2 probes, respectively).

Tyramide dye fluorophores (FITC, Cy3, Cy5: TSA plus, Perkin Elmer NEL760001KT) were used diluted appropriately in RNAscope TSA dilution buffer. Nuclei were labelled with RNAscope DAPI (ACDbio) for 30 seconds, or Hoechst (1/2000 in wash buffer for 10mins followed by two washes). Slides were mounted in Prolong Gold Antifade mounting medium (Thermofisher).

Positive and negative control slides were typically given the following fluorophore assignments: C1 –TSA-FITC; C2 – TSA-Cy3; C3 –TSA-Cy5. Due to weak FITC fluorescence and strong signal intensity of the 5’ dystrophin probe (as compared with the 3’ probe), many combinations of fluorophore posed either poor probe comparability or signal bleed-through (particularly those using TX2 filter cubes). Our preferred final fluorophore combination for dystrophin was therefore TSA-Cy3 and TSA-Cy5 (for C1 5’ and C2 3’, or switched as indicated). Used with the N3 and Y5 filter cubes, this combination of fluorophores had no signal overlap and allowed selection of modest exposure times without bleed-through. TSA dyes were used at the following dilutions: TSA-FITC, 1/500; TSA-Cy3, 1/1500; TSA-Cy5, 1/750.

### Imaging

Individual images were collected using either a DM4000B or DMRA2 upright fluorescence microscope with samples illuminated using an EBQ100 light source and A4, L5, N3 and Y5* filter cubes (Leica Microsystems) and an AxioCam MRm monochrome camera controlled through Axiovision software version 4.8.2 (Carl Zeiss Ltd). For all analysis, images were captured via 20x objectives (20x HC PL FLUOTAR PH2, NA=0.5) which readily enabled identification of discrete foci corresponding to individual transcripts, while retaining sufficient axial resolution to allow all elements of the tissue section to remain well-focussed. Multiple images (9–30) were collected per muscle section, selected randomly (avoiding section tears and tendinous regions). For particle counts, exposure times were calculated to allow clear detection of small foci in all channels (typically 600-1500msec). For fluorescence intensity (3-5 images per muscle section), multiple exposure times were collected (100-1000msec) to minimize risk of signal saturation. For whole section alignments, images were collected at 5x and stitched using the pairwise-stitching algorithm of Preibisch *et al* (29). After analysis, images shown in figures were adjusted for clarity using the window/level tool (imageJ).

### Image analysis

Collected images (as tiff format files) were analysed using ImageJ, using automated macros (all requisite.ijm files can be provided on request). In brief, images were separated into Cy5, Cy3 and DAPI channels: for counts of nuclei, nuclear stain was thresholded automatically using either Huang or Yen’s algorithm (30) and given a single binary watershed pass to separate adjacent nuclei. Cy3 and Cy5 channels were thresholded to eliminate background/non-specific staining and foci were counted using the ‘Analyze Particles’ tool. Fluorescence microscopy cannot reveal the true size of fluorescent particles (instead resolving a point-spread function (31)) but apparent size allows discriminatory analysis: apparent particle areas were converted to square microns using the appropriate multiplier for the magnification and image resolution. For nuclear/extranuclear particle analysis, the DAPI channel was thresholded as above and applied as a mask to the Cy3/Cy5 channels (eliminating nuclear-localized signal) or applied as an inverted mask (eliminating non-nuclear signal) prior to thresholding and particle counting. Nearest neighbour analysis used extranuclear foci only (distributions for random points were obtained from the average of ~1000 virtual fields of randomly-generated points of specified total number). Due to stark differences in 5’ signal intensity, 5’ probe fluorescence intensity analysis used images (3-5 per animal) collected at multiple exposure times (see above): regions of interest (sharply-defined small foci, broad nuclear probe signals, and background regions; 30-50 per image, 10-15 for background) were determined from the highest exposure, then quantified at lower exposure times using only non-saturated signal.

### Calculations

Mean values of transcripts per field were used to calculate total transcript numbers via the following calculations: 1388×1040 pixel imaging field at 20× (~2 pixels per micron) = 360880μm^2^ at a section thickness of 8um, thus 2889040μm^3^ (2.9×10^−3^mm^3^). At a muscle density of 1.06g.ml^−1^ (32), a single imaging field thus represents ~3μg of tissue. Typical RNA yields per mg of muscle tissue are 0.25-0.5μg, thus a single 20× imaging field can be considered to contain 0.75-1.5ng of total RNA.

#### Calculation of transcript half-lives

Half-life calculations used the method provided by Tennyson and colleagues (16). At steady-state, the ratio of 5’ sequence (nascent and mature dp427) to 3’ sequence (mature only) represents the balance of transcription time to mean lifetime.

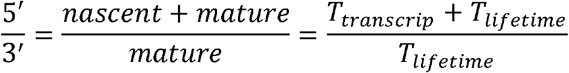

The half-life of a given species is related to the mean lifetime by the equation T_lifetime_ = T_1/2_/0.693.

Our 5’ probe sequence ends at exon 10 and the 3’ at exon 74 (1550kbp between the two) while our qPCR regions (see below) were designed to the exon 1:2 and exon 62:63 boundaries (1950kbp); assuming 16 hour constant-rate transcription for full-length dp427 (21) this gives an approximate transcription time between 5’ and 3’ of ~11 hours for RNAscope probe regions and 13.5 hours for qPCR amplicon regions.

### RNA isolation and qPCR

cDNAs used for qPCR (male quadriceps muscle, 9 WT, 9 *mdx*) were archive samples from our collection. Isolation and preparation methods were as described previously (33, 34): RNA was isolated from muscle powder (homogenized under liquid nitrogen) using TRIzol reagent (Invitrogen) with inclusion of an additional chloroform extraction (1:1) after phase separation. RNA was assessed to determine yield and purity (nanodrop ND1000) and used to prepare cDNA via RTnanoscript2 (Primerdesign). Triplicate qPCRs used 10μl volumes (~8ng cDNA per well assuming 1:1 conversion) in a CFX384 lightcycler using PrecisionPLUS SYBR green qPCR mastermix (Primerdesign), with a melt curve included as standard. Primers to *ACTB, RPL13a, CSNK2A2* and *AP3D1* (reference genes determined previously (33)) were taken from the geNorm and geNorm PLUS kits (Primerdesign). Primers to dystrophin dp427 and dp71 were designed using primer3 software (http://primer3.ut.ee/) and all span one or more introns (exons 62 and 63 are very short −61 and 62bp respectively-thus the reverse primer for this amplicon lies in exon 64, consequently spanning intron 62 and 63). All primer sets gave PCR efficiencies of 90-105% and all produced a single amplicon. Absolute quantification of transcript numbers was performed via standard curve (dilution series of purified, quantified PCR products from 10^Λ^7 molecules per well down to 10^Λ^−2), allowing accurate measurement of transcript numbers down to ~10-100 molecules.well^−1^ (typical values from muscle cDNAs were 500-50000 transcripts.well^−1^).

Dp427m Exon1 Fwd 5’-TCTCATCGTACCTAAGCCTCC-3’

Dp427 Exon2 Rev 5’-GAGGCGTTTTCCATCCTGC-3’

Dp427 exon44 Fwd 5’-TGGCTGAATGAAGTTGAACAGT-3’

Dp427 exon45 Rev 5’-CCGCAGACTCAAGCTTCCTA-3’

Dp427 exon62 Fwd 5’-AGCCATCTCACCAAACAAAGT-3‘

Dp427 exon64 Rev 5’-ACGCGGAGAACCTGACATTA-3’

Dp71 exon1 Fwd 5‘-GTGAAACCCTTACAACCATGAG-3’

Dp71 exon2 Rev 5’-CTTCTGGAGCCTTCTGAGC-3’

### Statistical analysis

Statistical analyses of nuclear counts and fluorescent foci counts used two-tailed Mann-Whitney U tests, while correlation analysis (5’ vs 3’ foci, total foci vs SDH or nuclear count) used Pearson r or Spearman’s rho as appropriate (indicated). Significance was set at P<0.05. All statistical analyses were conducted using Graphpad Prism 7.

## Results

### *In situ* hybridization reveals 5’ and 3’ sequences of dp427 mRNA show non-identical distributions, and allows detection of nascent and mature transcripts

RNAscope *in-situ* labelling allows resolution of individual mRNA molecules, particularly when used with low-abundance transcripts (24), and the expected staining pattern is that of small, punctate foci with an apparent diameter of 1-1.5μm distributed throughout the cell interior. Muscle labelled with positive control probes (to Polr2a, Ppib and UBC: very low, low and high-expression in skeletal muscle, respectively (35)) behaved consistently with these expectations (Supplementary Figure S2 A and B). Negative control probes (bacterial DapB in all channels) produced no foci (Supplementary Figure S2 C and D), though highly-diffuse, weakly-stained signal was found within small patches of cells in all dystrophic tissues, with all fluorophores (also observed in dystrophic positive control slides, but not in healthy muscle: most likely aberrant tyramide dye deposition by insufficiently-quenched endogenous peroxidase activity within the infiltrating macrophages common to dystrophic tissue). In contrast, muscle labelled with RNAscope probes to dystrophin 5’ and 3’ sequence revealed a striking and highly probe-specific distribution of signal. In healthy muscle (Figure 3) 5’ probe signal fell into two separable categories: the expected population of small, punctate foci distributed along the entire body of the myofiber (with the highest concentrations found closest to the sarcolemma), but also a second, less-numerous population of broad, heterogeneous and high-intensity signals localized exclusively to specific nuclei. The intensity of this latter population was such that these large 5’ foci were readily detectable even at low magnification. 3’ probe signal displayed essentially only the former punctate signal, both within nuclei and without; notably, these foci were typically found adjacent to 5’ foci. These distributions were consistent throughout the entire muscle section, both in transverse and longitudinal fibres, and reciprocal patterns were observed if the fluorophores chosen were exchanged (i.e. signal intensity and localization is a property of the probes alone: Supplementary Figure S3).

**Figure 3:**
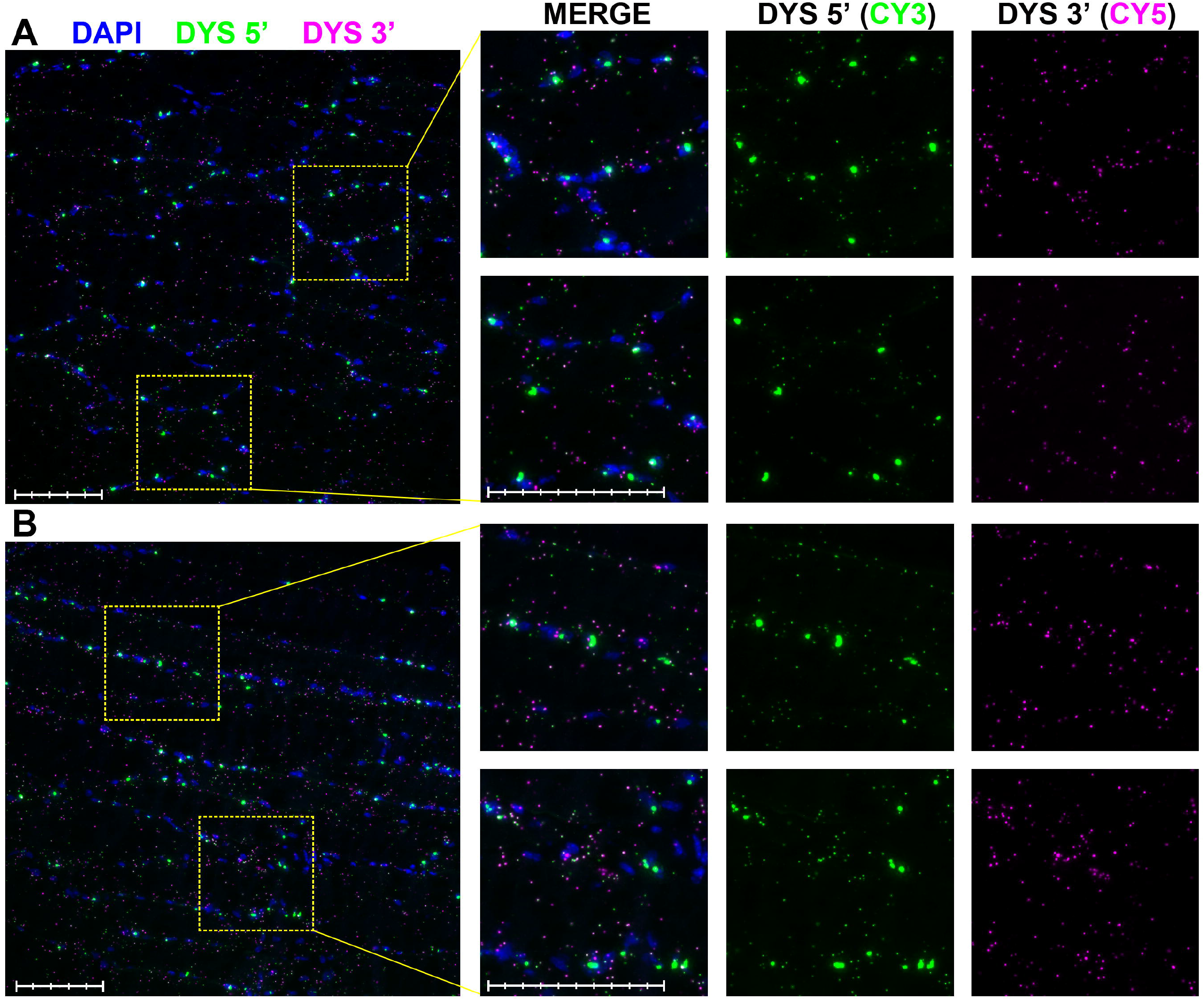
RNAscope labelling of dp427 5’ and 3’ in healthy mouse muscle. RNAscope probe labelling of 40-week old WT quadriceps muscle. Probe to dp427 5’ (Cy3: green) resolves both small sarcoplasmic foci and large nuclear foci, while 3’ probe (Cy5: magenta) shows small foci only. Foci are readily detected both in transverse (A) and longitudinal (B) section. Scale bars: 100μm (main panel subdivisions: 20μm; magnified panel subdivisions: 10μm).

Given co-transcriptional splicing (21), target sequence for the 5’ probe should emerge and persist early in transcription, while 3’ sequence should appear only shortly before mRNA completion and nuclear export. Intense nuclear staining for 5’ sequence (but not 3’) implies not only that nascent dp427 transcripts can readily be detected by our approach, but that multiple nascent transcripts are present within any given nucleus. The smaller foci (both 5’ and 3’) found outside nuclei would thus represent dual-labelling of mature dp427 transcripts. Supporting this, labelling of dystrophic *mdx* muscle with our probes (Figure 4) revealed key differences to healthy muscle: prominent 5’ nuclear foci were readily detected, but smaller punctate foci from both probe sets were much less prevalent and almost exclusively found within (or adjacent to) nuclei.

**Figure 4:**
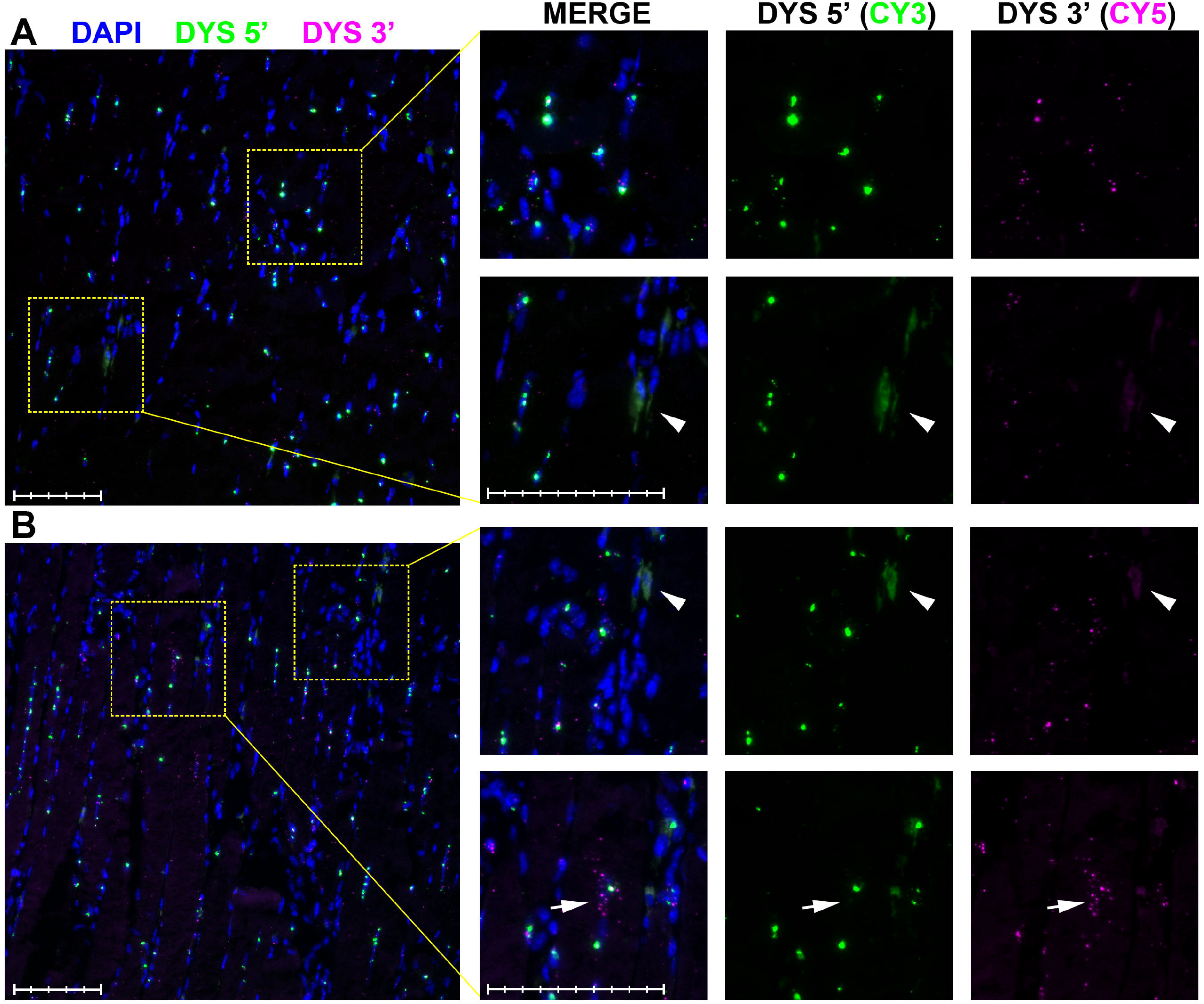
RNAscope labelling of dp427 5’ and 3’ in dystrophic mouse muscle. RNAscope probe labelling of 40-week old *mdx* quadriceps muscle. Probe to dp427 5’ (Cy3: green) resolves large nuclear foci but numbers of small foci are greatly reduced. 3’ probe (Cy5: magenta) reveals small foci only within or adjacent to nuclei. Nuclear foci are readily detected both in transverse (A) and longitudinal section. Rarely, cells with many 3’ foci not clearly associated with 5’ nuclear foci are observed (Arrows, lower panels). As with positive and negative control probes, apparently cell-restricted regions of nonspecific staining are frequently observed (Arrowheads, middle panels), likely corresponding to peroxidase-rich macrophage/neutrophil infiltration. Scale bars: 100¼m (main panel subdivisions: 20¼m; magnified panel subdivisions: 10¼m).

### RNAscope labelling of dp427 mRNA permits transcript quantification at subcellular levels

The well-defined foci produced by RNAscope labelling are amenable to semi-automated image analysis approaches, allowing measurement of both particle number and apparent area. As shown in Figure 5A and Table 1, healthy muscle particle counts were significantly higher than those for dystrophic muscle for both 5’ and 3’ probes. Per-image variation was however notable even within a given muscle section (Supplementary Figure S4A) and we speculated this might correlate with fibre type or myofibre nuclear number. Alignment of images with SDH-stained serial sections revealed a correlation between particle count and SDH activity (i.e. oxidative fibres might express higher levels of dp427) however these regions were also host to greater numbers of nuclei, and particle counts correlated better with nuclear number than with SDH (Supplementary Figure S4B-D), suggesting this is the principal source of within-section variation. Normalizing counts to nuclei would bias against dystrophic muscle (nuclear counts are significantly higher in this tissue: Supplementary Figure S4E), thus all analyses below used non-normalized, per-image counts.

**Figure 5:**
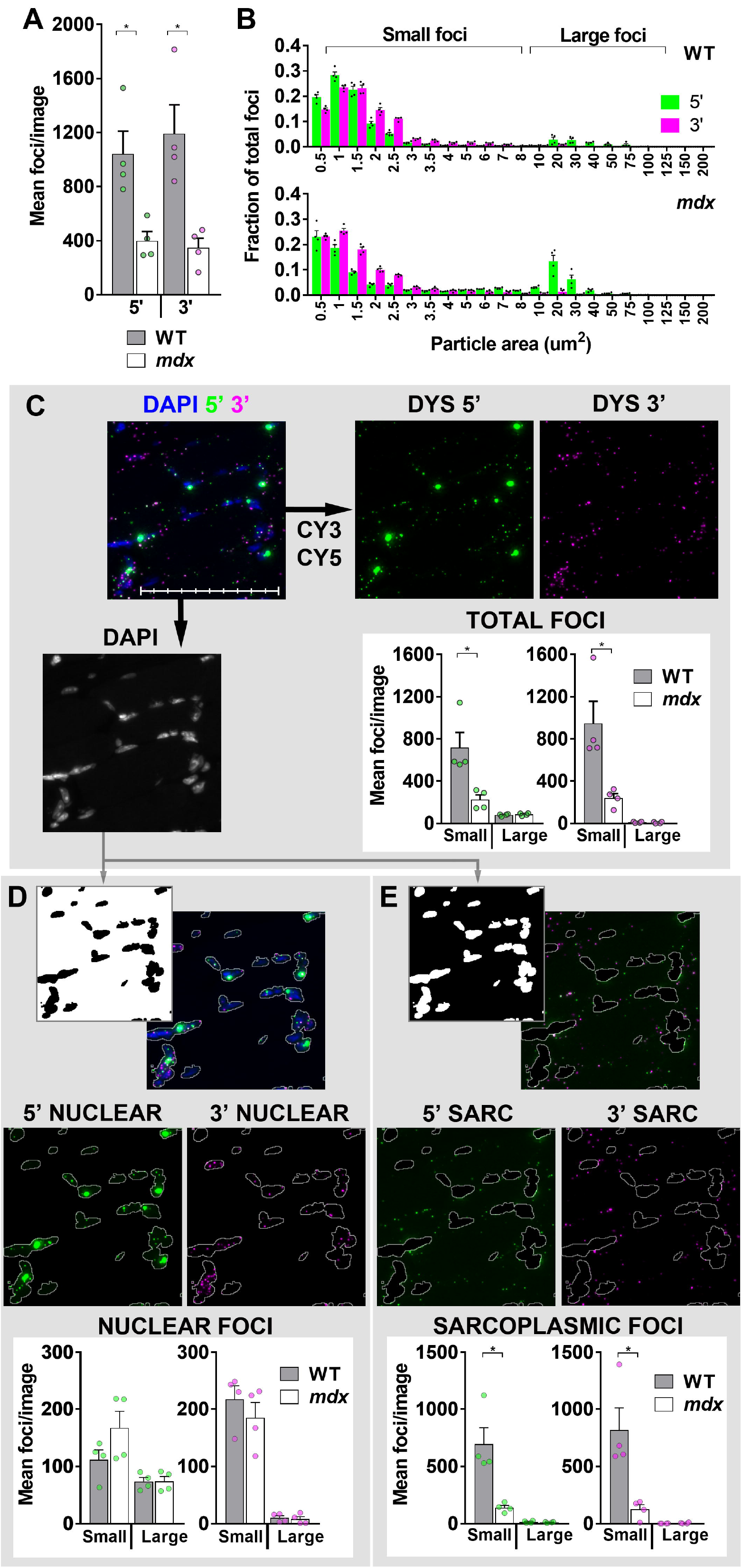
Particle analysis. (A) Total particle counts + SEM (N=4 per genotype) for 5’ and 3’ probes in healthy (WT) and dystrophic *(mdx)* muscle: values for each animal are shown. (B) Particle size distributions in WT and *mdx* muscle as fraction of total (per image) particle number: brackets used to define small/large foci are shown. (C) Segregation by small/large foci (as indicated) of total 5’ and 3’ signals, or signals after masking by nuclear stain (DAPI) to give nuclear (D –all sarcoplasmic foci removed) or sarcoplasmic (E –all nuclear foci removed) counts. Note that large foci are found only within nuclei, and only small sarcoplasmic foci (both 5’ and 3’) are significantly reduced in dystrophic muscle. Individual mean values per animal are shown as dots (∘: green = 5’; magenta = 3’) for all counts, or as dots (•) for size distributions. Each mean represents the average of 9-30 images (same panel of images used for all analyses). * = P<0.05, Mann-Whitney U test. Scale bar: 100μm (subdivisions: 10μm).

**Table 1:**
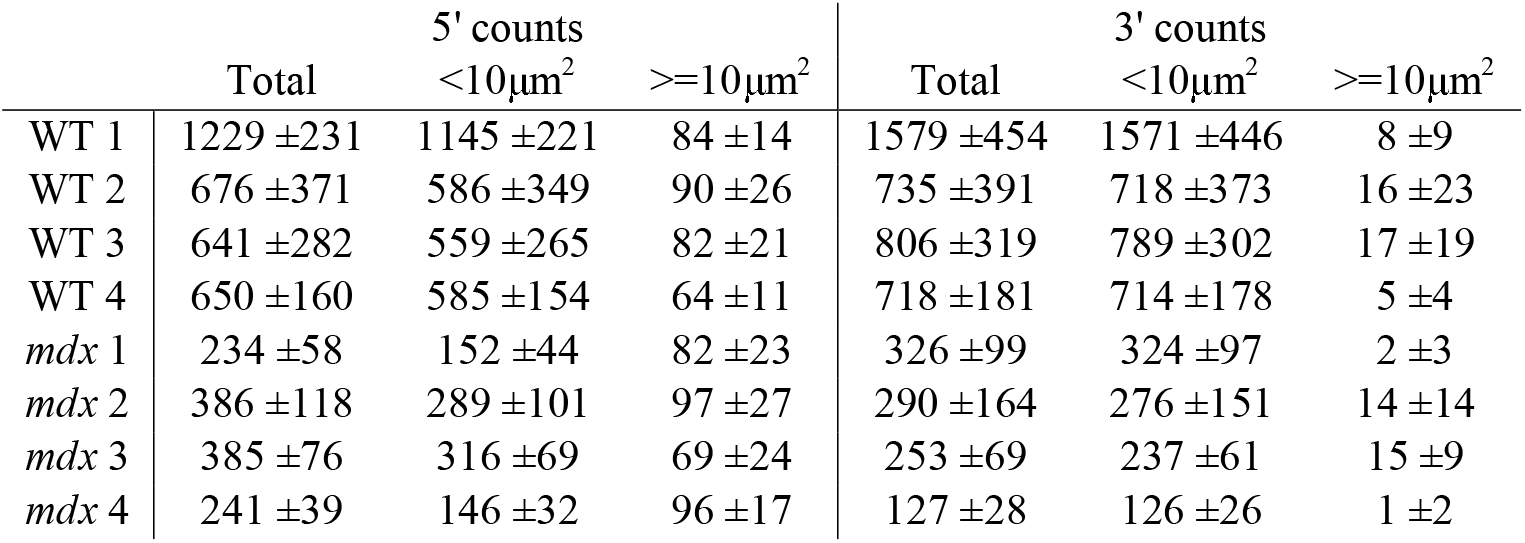
Mean per-image 5’ and 3’ particle counts for the eight quadriceps muscles used in this study. All counts expressed as total, or subdivided by size (all foci 0.5-10μm^2^ were classed as ‘small’, all those >10μm^2^ as ‘large’). Values shown are means ± standard deviations derived from 9-30 separate 20x images per muscle. Total and ‘small’ counts were significantly greater in WT muscle than dystrophic, while ‘large’ counts were not (P<0.05, Mann-Whitney U test). All mean values are shown graphically in figure 5A and C.

Despite marked differences in total count, healthy and dystrophic muscle exhibited similar distributions of apparent particle size (Figure 5B): signal from the 3’ probe was near-exclusively comprised of foci 1-2μm^2^ in area while 5’ probe signal could instead be segregated into two discrete populations, one of small foci similar to 3’ probe, and a second, larger and more heterogeneous population spanning 10-100μm^2^. We designated particles with areas between 0.5 and 10μm^2^ as ‘small foci’ (95% of this population fell between 1-3μm^2^) and those of 10-100μm^2^ as ‘large foci’. Segregated by these criteria (Figure 5C and Table 1), counts of large 5’ foci were comparable between healthy and dystrophic muscle: differences in 5’ particle number were entirely due to smaller foci. The unique multinucleate structure of myofibres permitted further segregation by subcellular distribution (nuclear or sarcoplasmic, Figure 5D and E), refining this analysis and confirming that large 5’ fluorophore deposits were indeed restricted to nuclei, while smaller foci (5’ and 3’) were found in both nuclear and sarcoplasmic compartments. Crucially, only the sarcoplasmic fractions of these small foci had reduced counts in dystrophic muscle: numbers of small foci within nuclei showed no reduction.

600-1400 sarcoplasmic foci were found in every healthy muscle image (10-20% within nuclei) compared with 100-300 in dystrophic muscle (50-60% nuclear). Large 5’ nuclear foci were present in lower numbers (50-100 per image in both healthy and dystrophic muscle), comprising only 6-12% percent of particles in healthy muscle (20-30% of nuclei) but accounting for 30-50% of total dystrophic counts (10-15% of nuclei due to the higher per-image nuclear count: Supplementary Figure S4E).

### 5’ and 3’ probes co-localize and reveal mature dp427 mRNAs within dystrophin positive revertant fibres

Extranuclear 5’ foci tended to be found close to 3’ foci in healthy muscle, and analysis also revealed a strong per-image correlation between 5’ and 3’ counts in this tissue, especially when restricted to small foci only: Pearson correlations in this fraction ranged from 0.91-0.97 (all P<0.0001), contrasted with 0.41-0.85 (P values from non-significant to <0.0001) for dystrophic muscle (Supplementary Figure S5 A and B). We measured the extent of probe co-localization in sarcoplasmic foci using a nearest-neighbour approach (determining minimum distance between extranuclear signals of opposing probes, compared with that found in an equivalent number of randomly distributed points). For large numbers of points even random distributions give the appearance of co-localization (with 1000 random points of each probe, 50% of foci can be paired with an opposing probe focus within a distance of 8μm or less, and within 20μm, 97% of foci are paired: see supplementary Figure S5 C and D), but numbers of foci paired within a micron are still low for random distributions (<1%). As shown in Figure 6, in healthy muscle almost 40% of all foci can be paired within this distance (whether 5’ to 3’ or the reverse), and of the remaining foci, minimum distances below 8μm are over-represented: pairings below 4μm were more common than would be expected even for 1400 random points (double the mean number of observed WT sarcoplasmic foci: see Supplementary Figure S5E). Conversely, the distribution of minimum distances in dystrophic muscle was more consistent with random chance, suggesting that the majority of sarcoplasmic foci (already low in number at ~100-200 per image) found in this tissue do not represent mature dp427 mRNAs (30-40% of foci were more than 30μm from an opposing focus, as seen with low numbers of random points: see Supplementary Figure S5F). Nevertheless, an increased fraction of paired distances below 8μm was observed, along with sub-micron colocalizations at levels approaching 25% in some images. Closer inspection showed these foci were not widely-distributed; instead occupying small fibre-restricted domains exhibiting local sarcoplasmic 5’/3’ counts comparable to healthy muscle. We conjectured these might correspond to corrected dp427 transcripts expressed within sporadic dystrophin-positive revertant fibres (36). Alignment with serial sections immunolabelled for dystrophin protein (Figure 7) showed that regions of dp427 probe labelling do indeed correspond to revertant fibres. RNAscope images collected only from regions aligning with revertant fibres showed higher mean sarcoplasmic 5’ counts than images collected randomly, though the increase was not uniform: some regions of revertant fibres showed no concomitant increase in probe labelling, implying that domains of dp427 mRNA might be more focal than those of dp427 protein.

**Figure 6:**
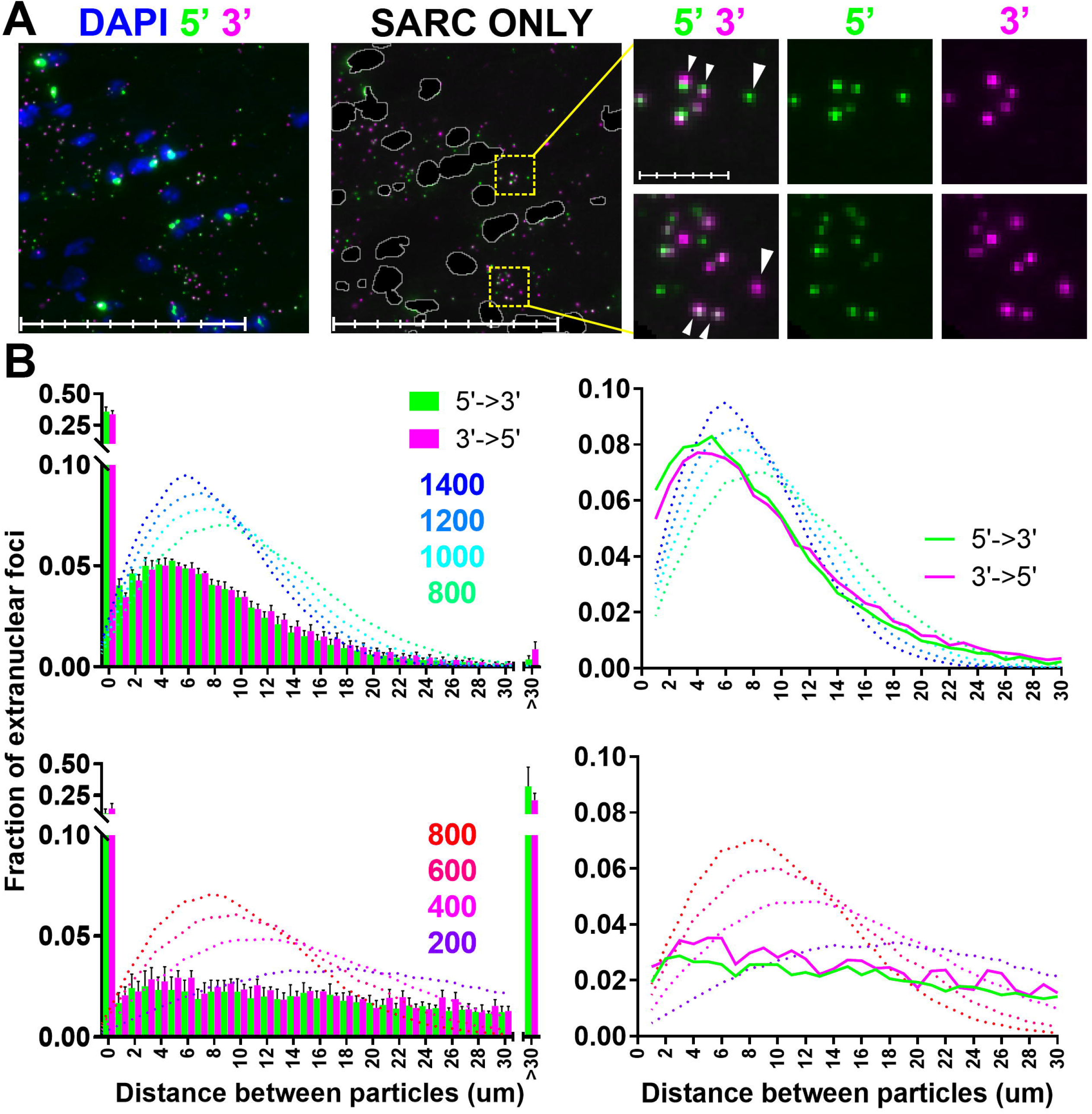
Co-localization of 5’ and 3’ signals. (A) 5’ and 3’ foci are found in close proximity. Images were masked to restrict analysis to sarcoplasmic signals only. Foci closer than 1μm are indicated (small arrowheads) though isolated foci of either probe were also observed (large arrowheads). Main panel scale bars: 100μm (subdivisions: 20μm); magnified panel scale bars: 10μm (subdivisions: 2 μm). (B) Distribution of distances between 5’ and 3’ sarcoplasmic foci. X-axis: distance between a given small 5’ sarcoplasmic signal and the nearest 3’ (or vice versa) in microns, Y-axis: fraction of total foci paired within that distance for WT (upper panels) and mdx (lower panels) quadriceps muscle. Left hand side: 5’-3’ distances are represented as green bars, 3’-5’ distances as magenta (bars = means+SEMs, N=4). 40% of WT and 10% of *mdx* foci co-localize within 1 micron while <1% of WT and >30% of *mdx* foci are more than 30μm from a focus of the opposite probe. Dotted traces represent expected mean distributions for 200-1400 random points (as indicated). Right hand side: fractional distribution of all distances >= 1 micron. Both WT (mean: ~800 foci per image) and mdx (mean: ~200) show a higher fraction of 5’-3’ pairings (or the reverse) below 8μm than would be expected.

**Figure 7:**
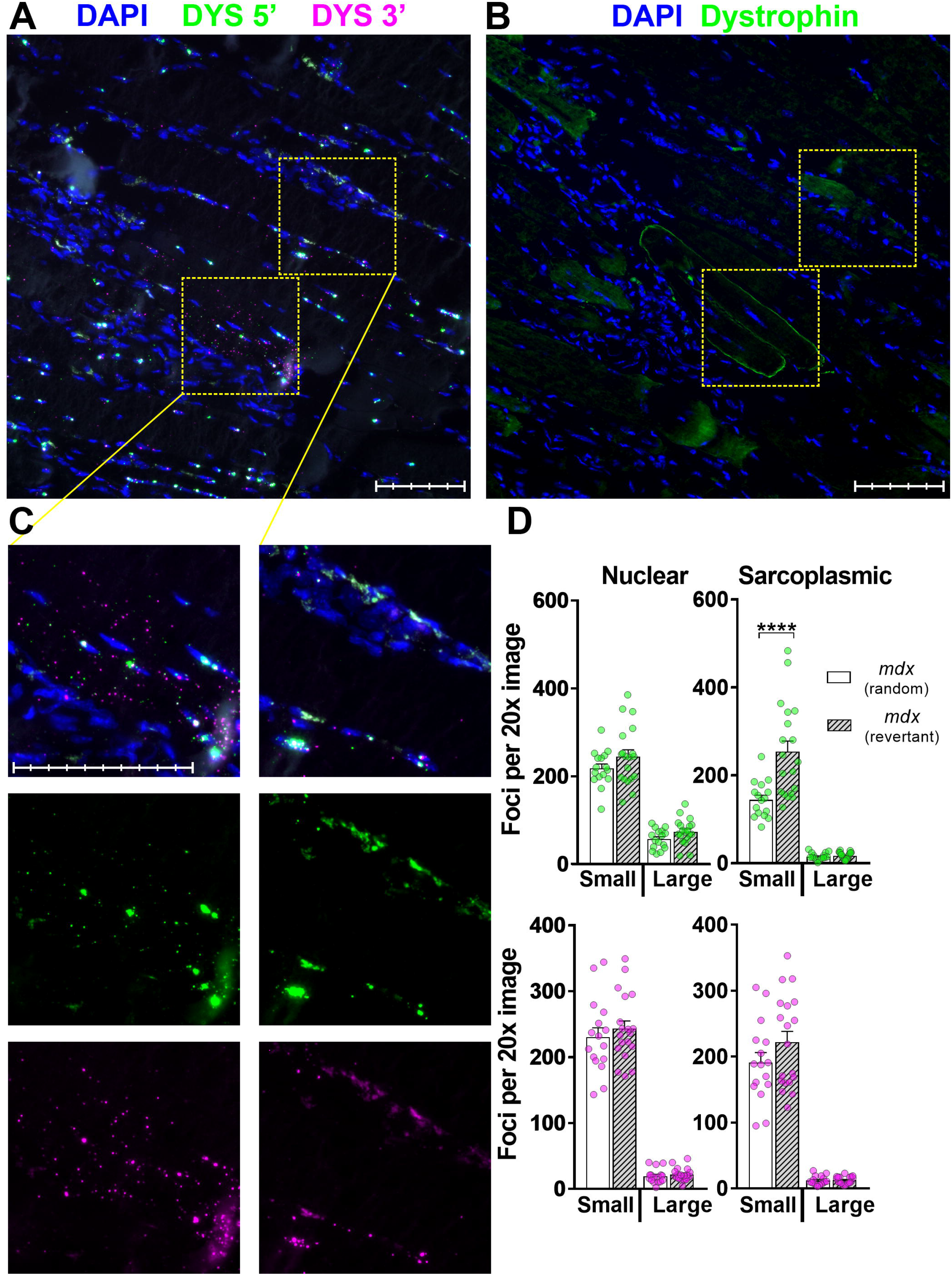
dp427 expression in revertant fibres. Aligned serial sections of *mdx* muscle labelled with RNAscope 5’/3’ probes (A) and dystrophin C-terminal antibody (B). RNAscope labelled regions corresponding to dystrophin-positive revertant fibres show large numbers of sarcoplasmic 5’ and 3’ foci (C, 1^st^ column) while those corresponding to adjacent dystrophin-negative fibres do not (C, 2^nd^ column). (D) nuclear and sarcoplasmic counts of 5’ and 3’ foci from a single RNAscope labelled dystrophic muscle using images collected at random or only from regions aligned with revertant fibres as indicated: revertant images show increased numbers of small sarcoplasmic 5’ foci (Each dot represents a single 20x image, P<0.0001, Mann-Whitney U test). Scale bars: 100μm (main panel subdivisions: 20μm; magnified panel subdivisions: 10μm).

Taken together, these observations support our interpretations: sharp, punctate foci outside the nuclear envelope –with 5’ and 3’ probe signals found in close apposition-are consistent with mature, exported dp427 mRNA molecules. Conversely, strong but less well-defined fluorescence restricted to the 5’ probe and found only within the nuclear compartment is consistent with labelling of immature transcripts. Concomitantly, only mature, exported dp427 signal shows profound reduction in dystrophic muscle: dp427 mRNA produced by *mdx* mice carries a premature termination codon (PTC) in exon 23 (37) and is subject to nonsense-mediated decay (NMD) (38), a checkpoint that occurs only after nuclear export (39).

### Quantification of nascent and mature transcripts shows mature dp427 mRNA is short-lived

As large nuclear foci represent nascent mRNA, an additional source of quantitation is offered by fluorescence intensity: the necessity for separate fluorophores precludes direct comparison between 5’ and 3’ probes, but careful choice of exposure time (Supplementary Figure S6) renders the small and large populations of 5’ probe foci tractable to such approaches. As extranuclear foci likely represent single transcripts, comparing mean fluorescence intensity of small sarcoplasmic signals and large myonuclear foci allows numbers of nascent dp427 molecules to be estimated. Our analysis (Figure 8A) suggests the dynamics of dystrophin expression are complex: in healthy muscle most myonuclei contain 20-40 immature transcripts, though expression falls over a considerable range: rare nuclei apparently have only 1-2 transcripts while others appear to contain 60 or more (though some of these higher values may reflect closely-apposed nuclei being counted together). Dystrophic myonuclei exhibit a similar spread of values, but expression as a whole is markedly reduced, with mean values suggesting only 10-12 nascent transcripts per nucleus.

**Figure 8:**
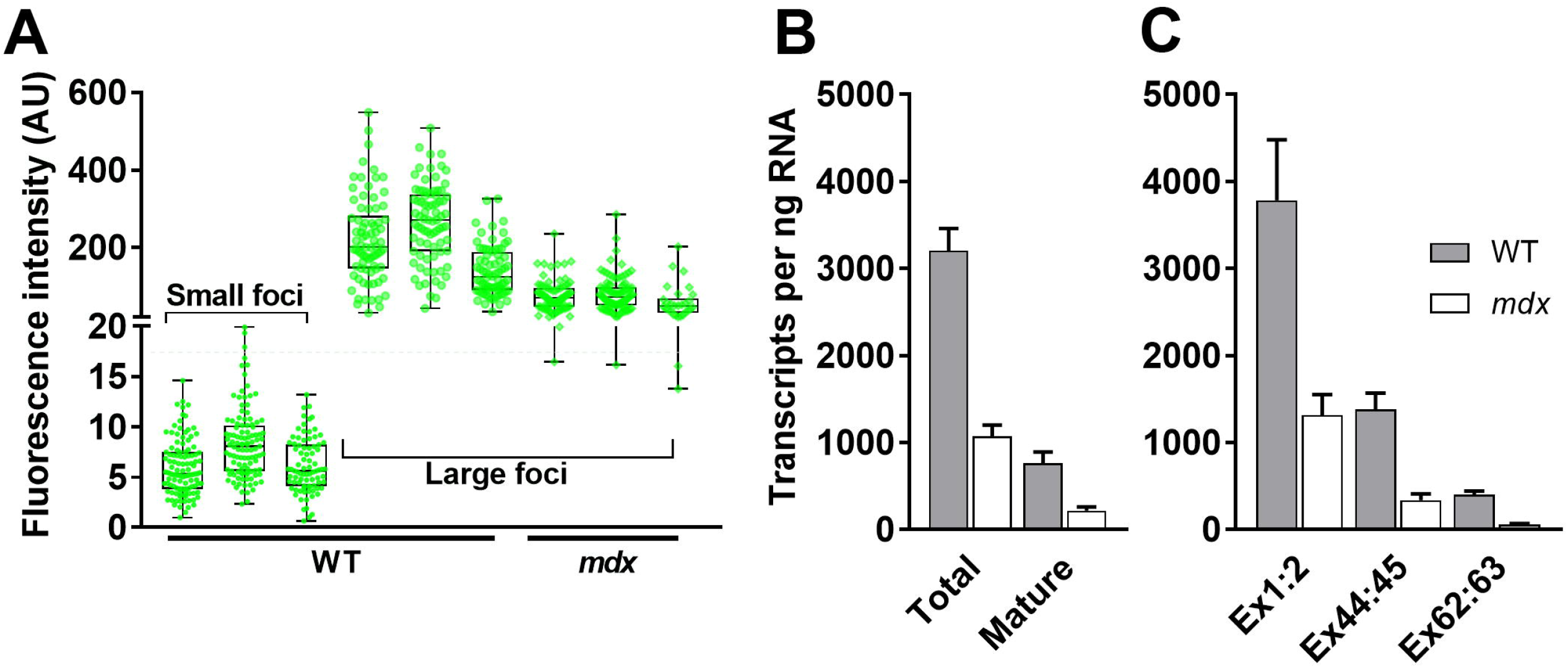
Nascent and mature dp427 transcripts. (A) Fluorescence intensity values from representative samplings of well-focussed non-saturated foci from small sarcoplasmic and large nuclear 5’ probe populations. Values obtained from three WT and three *mdx* mice (large foci only from *mdx* due to minimal numbers of small foci), using 3 images (60-120 particles in total) per individual. Values for small foci fall over a relatively narrow range (~5-8 AU) and can thus be used to estimate 5’ probe content of more heterogeneous large nuclear deposits (typically 100-400 WT, 50-150 *mdx)*. AU = arbitrary units. (B) Estimated numbers of dp427 transcripts in healthy and dystrophic mice by RNAscope counts: total transcripts (mean estimate of 5’ transcripts per nucleus multiplied by number of large nuclear foci, plus all mature foci) vs mature transcripts only, per 20x imaging field (equivalent to ~1ng RNA –see methods); (C) qPCR values: absolute transcript numbers per ng RNA as determined by qPCR to specific dystrophin dp427m exons as indicated, compared against standard curves. Data normalized to the geometric mean of APD3D1, ACTB, CNSK2A2 and RPL13a. All data shown as means + SEM (RNAscope N=4, qPCR N=9).

This data, combined with the counts of mature dp427 mRNA molecules above, allows estimation not only of total dystrophin expression, but also of the distribution of such expression between mature mRNA molecules and transcripts in progress (Figure 8B). Immature transcripts naturally comprise the bulk of signal in dystrophic muscle, but even within healthy muscle, prominent nuclear 5’ foci represent ~10% of the total 5’ signals: with each dp427-expressing nucleus host to 20-40 immature transcripts, the implication is that under normal healthy conditions nascent transcripts account for 60-80% of total dp427 mRNA.

We corroborated these estimates by quantitative RT-PCR (qPCR) targeted to specific 5’ or 3’ regions of the dystrophin gene: using primers spanning the exon:exon junctions at 1-2, 44-45 and 62-63, and quantifying absolute transcript numbers, we show that numbers of detected transcripts decline as expected as the target sequences approach the 3’ terminus, but in dystrophic muscle even 5’ signal is reduced, with transcripts containing exons 62-63 being reduced yet further (to levels approaching zero). As shown (Figure 8C) qPCR-derived transcript numbers also agreed closely with those obtained via RNAscope both for healthy and dystrophic samples, and confirmed that even in healthy muscle the bulk of dp427 is immature. This marked ~3-4-fold bias toward incomplete transcripts, combined with the estimated 16-hour transcription time for a single dp427 mRNA, suggests that once completed and exported, a mature dystrophin transcript has an average lifespan of only ~3.5 hours (see methods), substantially shorter than the time invested in its synthesis.

### RNAscope in situ hybridization permits quantification of dystrophin induction via exon skipping

To investigate whether dp427 probes could be used to assess therapeutic efficacy, we examined muscle treated with an exon-skipping agent. The *mdx* mouse carries a PTC in exon 23 of dp427: this exon lies inframe with those either side, and can thus be ‘skipped’ (omitted from the transcript during splicing): the resulting mRNA gives rise to internally-truncated but functional dystrophin protein. PIP6a PPMO molecules targeted to the splice donor site of exon 23 have been used to induce high levels of dystrophin in the *mdx* mouse: as shown previously (27), 10 doses of 12.5mg.kg^−1^ PPMO at 2-weekly intervals results in widespread high-level dystrophin protein expression (average ~50% of WT levels, with individual animals ranging from 30-80%) while a lower dose (6mg.kg^−1^) results in substantially lower expression (average ~15% of WT, with similar variability *–D. Wells, manuscript in preparation)*. Using a single quadriceps muscle from each study (high and low dose), we visualized PPMO induction of dp427 mRNA using RNAscope probes. As shown in Figure 9, lower dose treated muscle was essentially indistinguishable from untreated *mdx* muscle, but sarcoplasmic counts of both 5’ and 3’ foci in muscle treated with a higher dose were comparable to WT counts (and had concomitant restoration of 5’-3’ co-localization not seen with lower dose treatment: compare Supplementary Figures S7A and B). In high dose treated muscle, the fraction of nuclei with large 5’ foci was also restored to WT levels (25-30%) and nuclear 5’ signal was similarly more intense, yet numbers of nuclei per field remained elevated (comparable with untreated *mdx*, see Supplementary Figure S7C), with the corollary that all nuclear counts (5’ small/large and 3’ small) were higher than those of WT muscle.

**Figure 9:**
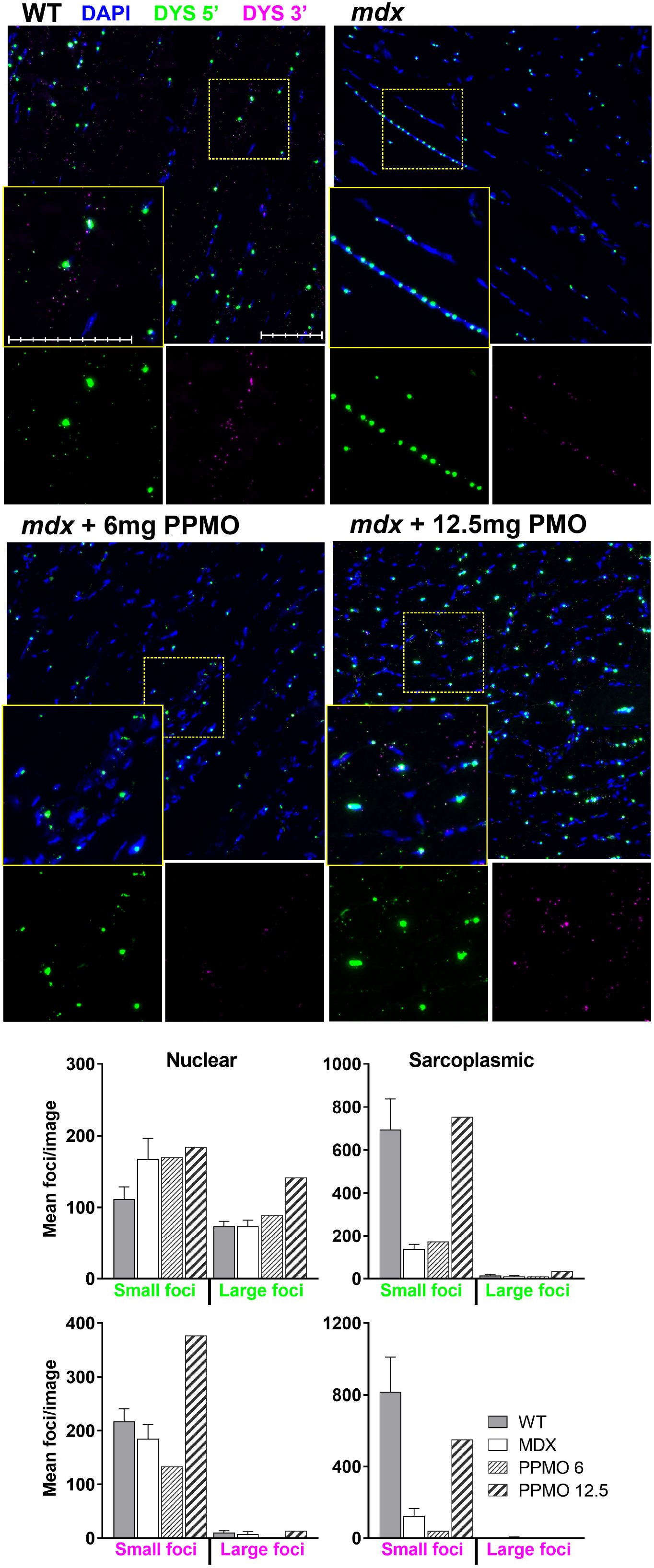
RNAscope detection of dp427 restoration following PPMO exon skipping. Top panels: RNAscope probes to dp427 5’ (Cy3: green) and 3’ (Cy5: magenta) in WT and mdx quadriceps muscle, or mdx muscle treated with PIP6a PPMO at 6mg.kg^−1^ or 12.5mg.kg^−1^ as indicated. High doses of Lower panels: numbers and distributions of 5’ and 3’ foci in PPMO treated muscles. WT and untreated mdx values are from figure 5. Treatment with 12.5mg.kg^−1^ restores small sarcoplasmic dystrophin counts to near-WT levels but also increases total nuclear counts of all foci.

## Discussion

Resolution of mRNA molecules by *in-situ* hybridization has historically faced a number of challenges: abundant transcripts can be detected relatively easily, but useful quantitative and spatial expression data is often lost in the consequent wash of strong signal. Conversely, low abundance transcripts might well lend themselves to more nuanced analysis, but cannot usually be detected with significant confidence to allow such analyses. Furthermore, most ISH methods are either radiometric or colorimetric, and thus cannot easily be multiplexed. The *in-situ* fluorescence amplification methodology of RNAscope ZZ-probes offers high sensitivity and multiplexing (24), adding a valuable tool for histological study of gene expression in health and disease. Although widely used for multiple mRNA species within a tissue, to our knowledge the data shown here represents the first use of this technology to visualize multiple probes within a single transcript. Even with ~1000 base hybridization sequences, the ~14,000 base dp427 transcript readily permits multiple probes to anneal; since expression is low, individual mature transcripts can readily be resolved as discrete entities. The enormous genomic DMD locus (~2.3Mbp, with 16-hour transcription time) further allows temporal metrics to be assessed, with strong 5’ probe labelling revealing nascent transcripts. Our work thus demonstrates a new use of RNAscope multiplexing that is particularly suited to long mRNAs transcribed from large genomic loci, allowing the identification of transcriptional subtleties that would otherwise remain intractable. Given that the DMD gene has key disease-relevance with many therapeutic approaches directed at the transcript level, our method is particularly suited to its study.

### Subcellular localization of mature dp427 mRNA

We show that individual mature dp427 mRNAs can be detected within the muscle sarcoplasm, binding both 5’ and 3’ probes in close proximity as expected. These mature transcripts are essentially absent in dystrophic muscle, yet comparatively high numbers (binding both probes) can still be found in rare patches that align with dystrophin immunoreactive revertant fibres. Previous in-situ studies of dystrophin dp427 expression have suggested that in healthy muscle this transcript preferentially localizes to the sarcolemma (17). Such a subcellular location is biologically plausible: the full-length dystrophin protein resides immediately beneath the sarcolemma, and this protein is moreover very large (427KDa in size) and carries multiple protein-binding domains. A single mRNA is relatively compact and can give rise to multiple protein molecules: targeting dp427 mRNA to the sarcolemma before translation would presumably be considerably more efficient than transporting protein post-translationally. The transcript-level resolution offered by our data suggests that this is largely correct: we have endeavoured to capture both longitudinal and transverse muscle sections, and while in both orientations mature dp427 molecules are clearly found throughout the myofiber interior, transcript concentration does indeed appear to be highest immediately beneath the sarcolemma.

### Behaviour and quantification of mature dp427 mRNA

The co-localization analysis permitted by our 5’/3’ labelling is revealing: under physiological conditions, most messenger RNA is expected to exhibit extensive secondary structure, and to be found as mRNP complexes with RNA binding proteins (40). This is especially true for mRNA molecules with large 5’ and 3’ UTRs (such as dystrophin), and indeed the 5’ and 3’ termini of such mRNAs are reported to be very closely apposed in most cases (potentially in the order of nanometre separations (41)). Efficient probe hybridization (and subsequent fluorophore labelling) requires conditions far from physiological, however, necessitating significant disruption of mRNP complexes and mRNA secondary structure (via protease treatment and lengthy incubations at supra-physiological temperatures in the presence of salts and detergents). One might therefore expect a significant percentage of transcripts to unfold to some extent, disrupting this colocalization. Despite this, in healthy muscle labelled with our dystrophin probes, ~40% of sarcoplasmic 5’ signals were found within a micron of a 3’ signal (and vice-versa), strongly suggesting that many transcripts are preserved in their native configuration (and that the physiological state of a mature dp427 mRNA is indeed a condensed complex with 5’ and 3’ ends in close proximity). The remaining foci were also typically paired within distances shorter than would be expected by chance, raising the possibility that these might represent labelling of unfolded transcripts. Length per base of single-stranded RNA remains contentious (42), and is further proposed to vary with salt concentration and pH (43), thus we cannot empirically state the maximum permissible separation of probes within a single dp427 mRNA. Measurements of ssDNA however report values of ~0.7nm per base (44), and using this value would put the length of a mature full-length dystrophin transcript at ~10μm, with the distance between our 5’ and 3’ probe binding sites just below 6μm (though ssRNA also adopts highly flexible, worm-like chain behaviour, thus mean 5’-3’ distance would be substantially shorter (45)). The size and mobility of the amplifier ‘trees’ generated by RNAscope ISH is not reported, and deposits of fluorophore (activated tyramides react with tyrosine residues in adjacent proteins) might also extend beyond these limits, but even restricting probe pairings to distances below 6μm would account for ~70% of the observed foci in WT muscle (~80% with a more generous 8μm). The remainder might also reflect limits of axial resolution (foci sufficiently out of focus as to be missed by particle analysis), or dp427 transcripts labelled with one probe but not the other (due to inhibitory secondary structure, masking by protein, or even cleavage of the molecule during sectioning). Our mature mRNA counts might consequently be an underestimate of true numbers, though our corroborating qPCR data suggests any such underestimate is modest. We note that counts of 3’ foci were typically higher than those for 5’, by a small but consistent amount (50-100), an observation partly explained by the greater ease with which signal from this probe can be collected (unlike 5’, all 3’ foci are of comparable size and intensity), and concomitantly by the fact that all 3’ foci represent single molecules, while 5’ foci in mature dp427 mRNAs lying adjacent to nuclei may be lost amidst strong nascent 5’ staining. 3’ labelling in the absence of adjacent 5’ might however also represent labelling of dp71: while not reported to be expressed in muscle fibres themselves, this transcript might nevertheless be found in blood vessel endothelia (46) or proliferating myoblasts (47), and is detected at low levels within whole muscle tissue homogenates, particularly those from young dystrophic muscle undergoing acute degeneration/regeneration (48). While our dystrophic muscle samples were taken from older mice (where pathology is established and consequently milder) we do observe rare nuclei with multiple nearby 3’ foci yet no 5’ labelling (see Figure 4B, lower panels). Dp71 expression measured by qPCR is low but detectable in quadriceps muscle (and comparable between healthy and dystrophic tissue): transcript numbers suggest this mRNA might account for ~50-70 additional 3’ foci per 20x image (Supplementary Figure S8).

### Behaviour and quantification of nascent dp427 mRNA

The genomic sequence of the dp427 region targeted by our 5’ probe (exons 2-10) begins ~190kb downstream from dp427m exon 1 and spans ~525kbp: assuming a constant, sequence-independent transcription rate of ~40bases.sec^−1^ with concurrent splicing (suggested by the 16-hour transcription time for a 2.3Mb gene, and supported by studies of other long genes (21, 22)), the first nucleotides of this sequence emerge ~1.3 hours after transcription commences, with the full 20ZZ probe-binding region being completed ~3.5 hours later and persisting within the nucleus for a further 11 hours (during which time additional dp427 transcripts can be initiated). In contrast, 3’ probe sequence (exons 64-75) begins 2140kb downstream of exon 1 and concludes a mere ~88kb later, only 28kb from the transcription termination site. Under the same assumptions, full 3’ probe sequence emerges 15.5 hours after initiation of transcription, less than 15 minutes before transcript completion (Figure 10A), and moreover subsequent export of completed mRNA from the nucleus is likely to be comparatively rapid (49) (especially compared to dp427 transcription time). RNAscope ISH provides only a snapshot of transcriptional state: we would thus expect to robustly capture high levels of 5’ probe signal within myonuclei expressing dp427, while nuclear 3’ probe foci should be smaller and found less frequently: exactly what we observe (notably, Smith *et al* reported similar findings in the nuclei of cultured myotubes using conventional fluorescent probes (25): while unable to resolve single transcripts, they reliably resolved the nuclear dp427 transcriptional locus).

**Figure 10:**
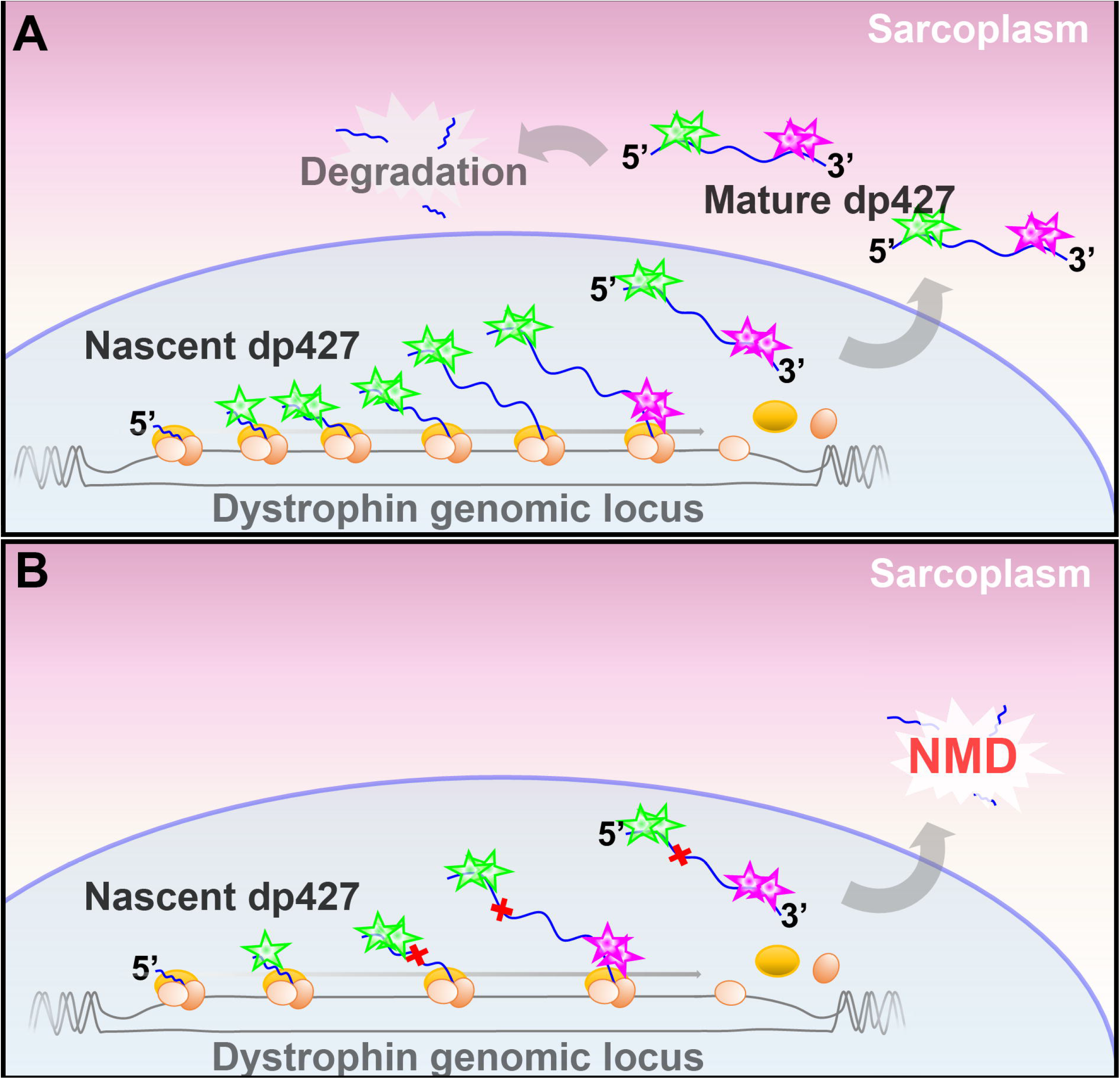
Dystrophin transcriptional dynamics revealed by RNAscope. In WT muscle myonuclei, RNA polymerases are loaded onto the transcriptional start site of the dystrophin locus faster than transcripts can be completed, resulting in a steady ‘train’ of polymerases and nascent transcripts, and leading to high nuclear concentrations of dp427 5’ sequence. Dp427 3’ sequence is completed only ~15 minutes before transcript completion and export, thus is present at much lower levels within the nucleus. Mature dp427 mRNAs, validated by the pioneer round of translation and carrying both 5’ and 3’ sequence, are found distributed throughout the sarcoplasm but are eventually degraded. In *mdx* muscle myonuclei, transcriptional initiation events are less frequent, leading to fewer nascent 5’ sequences within nuclei, and the presence of a PTC (red crosses) within the dp427 transcript leads to prompt post-export mRNA degradation via the NMD pathway.

A simple measurement permitted by this strong 5’ nuclear labelling is myonuclear number, a value otherwise difficult to estimate. Muscle is host to a variety of cell types, from blood-vessel endothelia and pericytes/mesoangioblasts to fibroblasts, adipocytes, and the dedicated muscle stem cell population of satellite cells (19, 20), but only myonuclei should robustly express full-length dystrophin. Our data thus suggest that in healthy muscle, myonuclei comprise 20-30% of the nuclei within a typical non-tendinous region of quadriceps muscle, while in dystrophic muscle this figure is 10-15% (commensurate with the higher fraction of non-myonuclear cell types within the dystrophic muscle environment).

A more nuanced quantification is the estimation of immature dp427 transcripts. As we show here, most dystrophin-expressing nuclei are host to 20-40 immature transcripts: if this represents steady-state occupancy, it implies active myonuclei produce one dp427 transcript every 20-30 minutes (though as ~5 hours are required to transcribe the full 5’ binding sequence, true nascent counts may be ~30% higher). Both our RNAscope and qPCR data suggest that mature dp427 mRNAs are in the clear minority: 60-80% of dp427 transcripts within healthy skeletal muscle are nascent. The surprising corollary is that despite taking 16 hours to produce, the half-life of mature dp427 is only ~2-3 hours (mean lifetime of 3-4 hours). Similar results were first reported in the late 90s by Tennyson and colleagues (16) using radiolabelled RT-PCR on samples of human skeletal muscle, and the values obtained were remarkably close to those shown here. These early studies obtained mean 5’ and 3’ totals of ~2500 and ~780 per ng RNA respectively (interestingly, with sample-to-sample variation similar to that found in our study), with concomitant derived half-life of ~3.5 hours. This early finding has garnered little mention since –even the authors themselves treated this result with scepticism-but our work here (showing similar values in a different species) strongly supports this remarkable result. Tennyson *et al* reported a markedly longer half-life of ~16 hours in cultured human foetal myotubes: this discrepancy was attributed to either differential turnover rates between cultured myotubes and mature muscle, a higher rate of premature termination in mature muscle, or an indication that mature skeletal muscle was not at transcriptional steady-state (samples were collected from human cadavers). Our data, obtained from mouse muscle flash-frozen 2-5 minutes post-mortem, would tend to argue against this latter hypothesis. Relatively high rates of premature termination are certainly plausible: at 2.3 million bases, the dystrophin gene likely tests the limits of RNA polymerase processivity (50), and some evidence exists for premature termination rates above predicted values (21), but such limits are a function of length, not maturity. It is thus unclear why such rates would differ markedly between myogenic cultures and mature muscle. Dp427 may simply experience a higher rate of turnover in mature muscle: the demand for dystrophin is likely to be higher in nascent myotubes where the contractile environment is being established *de novo* (unlike mature skeletal muscle where turnover of an existing dystrophin protein pool may be all that is required). Given the gene size, production of dystrophin transcripts could be considered innately wasteful (synthesis of a 2.3Mb pre-mRNA for a final transcript of only 14kb), but further, it seems remarkable that mature muscle myonuclei may indeed dedicate 16 hours to producing a molecule with a mean subsequent lifespan of only 4. Biological systems do however often favour expediency over efficiency, and dp427 transcripts comprise only a tiny fraction of myonuclear mRNAs (several thousand-fold lower than myosin heavy chain mRNA (16)) thus such waste is likely of little energetic consequence. Post-transcriptional control via mRNA stability in fact offers markedly greater utility than control of transcription itself: assuming the necessary transcription factors could be mobilized rapidly, any changes in expression based on transcriptional initiation would necessarily incur a 16-hour delay (and a second 16-hour delay to return to basal levels) and this presumes the dystrophin locus is not already at near-maximal transcriptional occupancy. In contrast, changes in mRNA stability can be achieved rapidly via phosphorylation (51), and simply halting degradation of mature dp427 transcripts (leaving transcription unchanged) would double mRNA levels in less than half that time. This wasteful but responsive approach might even be common among large genes: the human genome is host to a number of genes larger than 1Mb (>7 hour transcription times), many of which may need to alter expression over shorter timescales. Dynamic expression of dp427 would readily explain the high sample-to-sample variation reported here and previously (16), and control via mRNA stability would allow expression to fluctuate over a circadian timescale (as has been suggested (52)). It would be of great interest to examine the behaviour of our RNAscope probes in damaged (Cardiotoxin or BaCl_2_-treated), regenerating healthy muscle: if stability of transcripts is higher in nascent myotubes one might expect newly regenerated fibres to exhibit markedly higher levels of mature dp427 mRNA (something suggested by early ISH work (18)).

Tennyson *et al* also derived estimates of 5-10 nascent dystrophin mRNAs per muscle nucleus (16). Our data here refines this estimate: not all nuclei within a muscle express dystrophin (see above), but those that do are host to 20-40 nascent transcripts at a time. This argues in favour of a concerted commitment to dystrophin expression, and provides further support for a maximized production hypothesis. Rare nuclei nevertheless appear to have lower (2–10) numbers of transcripts, suggesting that dystrophin-expressing nuclei might not all be at steady-state expression levels at any given moment: we cannot rule out a sustained but ‘burst-like’ stochastic expression programme as is common to many eukaryotic genes (53) (transcription in muscle has been reported to exhibit ‘pulsed’ behaviour (54)). Transcriptional bursting could result in a number of ‘quiescent’ myonuclei that do not label with our 5’ probe (leading to underestimation of myonuclear fraction), though counts of dp427-expressing nuclei were remarkably consistent between animals, suggesting this is unlikely (Table 1 and figure 5D).

Dystrophic muscle reveals further nuances: while numbers of mature transcripts are reduced to almost zero as expected, the intensity of 5’ probe labelling within myonuclei is also reduced. Similar reductions in total message are seen when measurements are made by qPCR. While it would be easy to dismiss this latter observation as a reflection of the higher cell content and transcriptional diversity within dystrophic muscle (thus fewer total myonuclei), our RNAscope data shows that this reduction in expression extends to the myonuclei themselves. As noted, transcript degradation via NMD occurs only after transcription is complete (39), thus reduction in nuclear signal is best explained by a lower rate of transcriptional initiation (10-12 nascent transcripts per nucleus, thus one transcript initiated/completed every 60-90 minutes). It has been proposed that as a terminal marker of myogenesis, expression of dystrophin itself drives the myogenic programme to completion: low levels of dystrophin expressed earlier in differentiation help drive greater expression of dystrophin, further promoting terminal differentiation and producing a switch-like feedforward loop (55, 56). Dystrophic muscle, lacking this driver of terminal myofiber commitment, would thus be held in a permanent state of near-maturation, with concomitantly lower levels of dystrophin transcription and reduced 5’ nuclear staining intensity (Figure 10B).

Our dp427 probes were also capable of recognizing transcripts associated with revertant fibres: the presence of these sporadic dystrophin-positive fibres within dystrophic muscle has long been recognized (36, 57, 58), and variably explained. Prevalence of these fibres is influenced by the specific mutation carried (the PTC in exon 23 of the *mdx* mouse readily lends itself to revertant fibre formation, while the PTC in exon 53 of the *mdx^4cv^* mouse does not (59)), and revertant fibre number also tends to increase with age, clustering in a fashion that implies a clonal origin (60, 61) (skipped transcripts are found even in healthy muscle (62)).

Current hypotheses tend to favour a coordinated and somehow focally heritable alteration in splicing patterns rather than mutations at the genomic level (60): our data here do not allow these two proposals to be discerned, and indeed the high nuclear counts in dystrophic muscle makes assignation of revertant expression to a specific nucleus or nuclei essentially impossible, but our data do suggest that revertant fibres exhibit high (near-WT) levels of mature dp427 within relatively restricted domains (as might be expected if transcripts are short-lived). We also show that our probes can be used to assess the efficacy of exon skipping approaches, and that our data closely matches that reported previously via alternative methods while providing additional spatial metrics. *Mdx* mice treated with PIP6a peptide-conjugated skipping oligo at 6mg.kg^−1^ show low levels (2-10%) of exon skipping and consequently low induction of dystrophin protein (~15%), while treatment at 12.5mg.kg^−1^ gave skipping levels as high as 80%, widespread dystrophin protein induction, almost complete protection against eccentric exercise and restoration of muscle force generation (27). Our data suggests that assessment of skipping percentages via qPCR, especially for an exon nearer the 5’ end of the dp427 transcript (such as exon 23) might prove less informative than expected: values expressed as a ratio provide no information regarding absolute numbers, and while successfully skipped transcripts will certainly give rise to mature dp427 mRNAs (unlike unskipped, which will promptly be degraded), even in healthy muscle, mature mRNA represents only ~20% of total dp427; the majority of the dp427 measured (skipped or unskipped) will be in the form of nascent rather than mature transcripts. Conversely, labelling these muscles using our RNAscope probes does not permit assessment of skipping percentages, but instead allows both quantification of mature transcripts and estimation of total transcript numbers. While only a single muscle at each dose was examined, as we show here, 6mg.kg^−1^ treated muscle was indistinguishable from untreated *mdx*, while muscle treated with 12.5mg.kg^−1^ PPMO showed sarcoplasmic dp427 mRNA counts comparable to those of WT muscle. Our data suggests RNAscope *in-situ* methods can also reveal changes in transcriptional dynamics: counts of nuclei per field in 12.5mg.kg^−1^ PPMO treated muscle were comparable to untreated *mdx* while the fraction of nuclei with large 5’ foci was comparable to healthy muscle (i.e. total numbers were higher than WT), suggesting that even without 100% skipping efficiency, treated muscles might achieve physiological levels of mature dp427 by producing supra-physiological levels of nascent transcripts. Fluorescence intensity of nuclear 5’ foci was also increased over untreated *mdx*, implying that restoration of dystrophin contributes to the feed-forward mechanism, driving myofibre commitment and enhanced dp427 expression. This analysis raises a number of questions, however: in mice skipped transcripts are typically detectable within a week of systemic treatment, but a study following long-term response to exon skipping treatment in the *mdx* mouse (63) reported a 2-3 week lag between first detection of skipped transcripts and restoration of dystrophin protein, and further showed that both skipped dp427 transcripts and particularly dystrophin protein itself persisted for some weeks following the cessation of treatment. Levels of skipped transcripts and dystrophin protein in this study were low (~10% and 1-3% of WT respectively) but these data nevertheless suggest that the mean lifespan of dystrophin protein is of the order of days to weeks, and moreover that protein turnover is comparatively slow. It is hard to reconcile this with the mRNA dynamics revealed by our studies here (and those reported historically (16)): despite a lengthy transcription time, mRNA turnover of dp427 appears to be remarkably rapid. Tissue persistence of antisense oligonucleotides was observed to decline in line with falling skipping percentages (63) (a process ostensibly compatible with ongoing skipping of short-lived dp427 mRNAs), the high stability of the dystrophin protein and the fact that this protein represents only a relatively minor component of the muscle protein milieu (14) seems incompatible with such dynamic transcript behaviour. The implication is that in healthy muscle, dp427 mRNA molecules may be actively degraded despite having contributed essentially no dystrophin protein beyond the first post-export pioneering ribosomal proof-read.

If correct, this surprising finding might in fact be of considerable value to therapeutic approaches for DMD such as gene editing: if under healthy conditions many more transcripts are produced than are needed, in dystrophic conditions, beneficial levels of dystrophin protein might be restored with only modest levels of genomic correction (and studies suggest this may be the case (64–68)).

### Concluding remarks

The work shown here validates a novel use of RNAscope methodology, demonstrating the value of multiplex targeting within a single transcript, particularly with mRNAs transcribed from long genomic loci such as the dystrophin gene. The ability to visualize individual dystrophin transcripts –and individual regions of dystrophin transcripts-at subcellular resolution within tissue sections suggests a number of enticing applications, both those from a purely biological perspective, and those pertaining to human medicine. Our studies have focussed on dystrophin expression in skeletal muscle: expression in this tissue is comparatively well-understood; samples of healthy, dystrophic and exon-skipping treated quadriceps are readily obtained; and this tissue is of key disease relevance (indeed our approach can be used to assess therapeutic efficiency). There is no *a priori* reason why studies of dystrophin expression need be restricted to muscle however, and this latter consideration merits further clarification. The dystrophin gene is remarkable in that a single (admittedly enormous) gene gives rise to multiple mature mRNAs (and thus multiple protein isoforms) that are not splice variants, but 5’ truncated sequences arising from different internal promoters. Moreover, expression of these isoforms (full length dp427, dp260, dp140, dp116 and dp71) is regulated in a highly tissue-specific fashion, suggesting that these truncated proteins all serve key functional roles. The shared sequence identity both at mRNA and protein level makes resolution of individual isoforms challenging, and spatiotemporal resolution within a given tissue is essentially impossible by conventional approaches. The ability to bind ISH probes to specific sequences within the dystrophin gene in a multiplex fashion offers a means to achieve just such spatiotemporal resolution. We have used two probes here: one to 5’ sequence (exons 2-10), and one to 3’ (exons 63-74). As shown, full length dystrophin (dp427) binds both probes, and the fluorophore signals are found in very close proximity. Moreover, nuclei expressing full-length dystrophin are rendered obvious by virtue of the high levels of 5’ probe binding to nascent dp427 transcripts: expression of dp427 can be detected very clearly. A corollary of this is that robust 3’ probe signal in the absence of matching adjacent 5’ signal must therefore correspond to dp260, dp140 or dp71 (Figure 1): these probes potentially allow in-situ discrimination of different dystrophin mRNA isoforms. RNAscope offers multiplexing beyond 2 probes: 3 or even 4 probe channels are feasible. Given the potentially widespread expression profiles of dp140 and dp71 (4, 6, 47) (dp260 is believed to be largely retina-restricted (3)), a logical choice for a third probe would be one capable of distinguishing dp140 from dp71 (for example, binding to exons45-55). A triplet of probes (5’, middle, 3’) would allow extremely high resolution of multiple dystrophin isoforms: dp427 would show all 3 probes in close association, dp140 would show only 2, and dp71 would label with the 3’ probe alone. Such a probe set might allow the intricacies of systemic dystrophin expression during neonatal development to be resolved, or allow the unique distribution of dystrophin isoforms within the brain to be mapped down to subcellular levels. Indeed, while the current therapeutic focus is on skeletal muscle, the brain is also known to be affected by dystrophin mutations: if treatment of skeletal muscle continues along current encouraging lines, the brain might be the next major therapeutic target, and the *in situ* hybridization methodology described here offers a powerful tool to aid such investigations.

## Nonstandard Abbreviations

ISH: In situ hybridization
DMD: Duchenne muscular dystrophy
BMD: Becker muscular dystrophy
NMD: Nonsense-mediated decay
DAGC: Dystrophin-associated glycoprotein complex
nNOS: neuronal nitric oxide synthase
PPMO: PIP6a peptide-conjugated antisense morpholino oligonucleotide
FITC: Fluorescein isothiocyanate
Cy3/Cy5: Cyanine 3/5
SDH: Succinate dehydrogenase
PTC: Premature termination codon

## Acknowledgements

This work was funded by the Wellcome Trust [Grant number 101550/Z/13/Z].

The authors declare they have no conflicting interests.

This manuscript was approved by the RVC research office and assigned the following number: CSS_02001

## Author Contributions

J.C.W. Hildyard and R.J. Piercy designed research; J.C.W. Hildyard and F. Rawson performed research; J.C.W. Hildyard analysed data; D.J. Wells contributed tissue samples; J.C.W. Hildyard wrote the manuscript; F. Rawson, D.J. Wells and R.J. Piercy reviewed the manuscript.

## References

1. Muntoni, F., Torelli, S., and Ferlini, A. (2003) Dystrophin and mutations: one gene, several proteins, multiple phenotypes. Lancet Neurol 2, 731–740

2. Gorecki, D. C., Monaco, A. P., Derry, J. M., Walker, A. P., Barnard, E. A., and Barnard, P. J. (1992) Expression of four alternative dystrophin transcripts in brain regions regulated by different promoters. Human molecular genetics 1, 505–510

3. D’Souza, V. N., Nguyen, T. M., Morris, G. E., Karges, W., Pillers, D. A., and Ray, P. N. (1995) A novel dystrophin isoform is required for normal retinal electrophysiology. Human molecular genetics 4, 837–842

4. Lidov, H. G., Selig, S., and Kunkel, L. M. (1995) Dp140: a novel 140 kDa CNS transcript from the dystrophin locus. Human molecular genetics 4, 329–335

5. Byers, T. J., Lidov, H. G., and Kunkel, L. M. (1993) An alternative dystrophin transcript specific to peripheral nerve. Nature genetics 4, 77–81

6. Bar, S., Barnea, E., Levy, Z., Neuman, S., Yaffe, D., and Nudel, U. (1990) A novel product of the Duchenne muscular dystrophy gene which greatly differs from the known isoforms in its structure and tissue distribution. Biochem J 272, 557–560

7. Mandel, J. L. (1989) Dystrophin. The gene and its product. Nature 339, 584–586

8. Sadoulet-Puccio, H. M., and Kunkel, L. M. (1996) Dystrophin and its isoforms. Brain pathology (Zurich, Switzerland) 6, 25–35

9. Brenman, J. E., Chao, D. S., Xia, H., Aldape, K., and Bredt, D. S. (1995) Nitric oxide synthase complexed with dystrophin and absent from skeletal muscle sarcolemma in Duchenne muscular dystrophy. Cell 82, 743–752

10. Campbell, K. P., and Kahl, S. D. (1989) Association of dystrophin and an integral membrane glycoprotein. Nature 338, 259–262

11. Stark, A. E. (2015) Determinants of the incidence of Duchenne muscular dystrophy. Ann Transl Med 3, 287

12. Mah, J. K., Korngut, L., Dykeman, J., Day, L., Pringsheim, T., and Jette, N. (2014) A systematic review and meta-analysis on the epidemiology of Duchenne and Becker muscular dystrophy. Neuromuscular disorders: NMD 24, 482–491

13. Yiu, E. M., and Kornberg, A. J. (2015) Duchenne muscular dystrophy. J Paediatr Child Health 51, 759–764

14. Hoffman, E. P., Brown, R. H., Jr., and Kunkel, L. M. (1987) Dystrophin: the protein product of the Duchenne muscular dystrophy locus. Cell 51, 919–928

15. Muntoni, F., and Strong, P. N. (1989) Transcription of the dystrophin gene in Duchenne muscular dystrophy muscle. FEBS Lett 252, 95–98

16. Tennyson, C. N., Shi, Q., and Worton, R. G. (1996) Stability of the human dystrophin transcript in muscle. Nucleic acids research 24, 3059–3064

17. Mitsui, T., Kawai, H., Shano, M., Kawajiri, M., Kunishige, M., and Saito, S. (1997) Preferential Subsarcolemmal Localization of Dystrophin and β-dystroglycan mRNA in Human Skeletal Muscles. Journal of neuropathology and experimental neurology 56, 94–101

18. Scott, M. O., Sylvester, J. E., Heiman-Patterson, T., Shi, Y. J., Fieles, W., Stedman, H., Burghes, A., Ray, P., Worton, R., and Fischbeck, K. H. (1988) Duchenne muscular dystrophy gene expression in normal and diseased human muscle. Science (New York, N. Y.) 239, 1418–1420

19. Ceafalan, L. C., Popescu, B. O., and Hinescu, M. E. (2014) Cellular players in skeletal muscle regeneration. Biomed Res Int 2014, 957014

20. Tanabe, Y., Esaki, K., and Nomura, T. (1986) Skeletal muscle pathology in X chromosome-linked muscular dystrophy (mdx) mouse. Acta Neuropathol 69, 91–95

21. Tennyson, C. N., Klamut, H. J., and Worton, R. G. (1995) The human dystrophin gene requires 16 hours to be transcribed and is cotranscriptionally spliced. Nature genetics 9, 184–190

22. Singh, J., and Padgett, R. A. (2009) Rates of in situ transcription and splicing in large human genes. Nat Struct Mol Biol 16, 1128–1133

23. Gazzoli, I., Pulyakhina, I., Verwey, N. E., Ariyurek, Y., Laros, J. F., t Hoen, P. A., and Aartsma-Rus, A. (2016) Non-sequential and multi-step splicing of the dystrophin transcript. RNA Biol 13, 290–305

24. Wang, F., Flanagan, J., Su, N., Wang, L. C., Bui, S., Nielson, A., Wu, X., Vo, H. T., Ma, X. J., and Luo, Y. (2012) RNAscope: a novel in situ RNA analysis platform for formalin-fixed, paraffin-embedded tissues. J Mol Diagn 14, 22–29

25. Smith, K. P., Moen, P. T., Wydner, K. L., Coleman, J. R., and Lawrence, J. B. (1999) Processing of endogenous pre-mRNAs in association with SC-35 domains is gene specific. The Journal of cell biology 144, 617–629

26. Garcia, H. G., Tikhonov, M., Lin, A., and Gregor, T. (2013) Quantitative imaging of transcription in living Drosophila embryos links polymerase activity to patterning. Curr Biol 23, 2140–2145

27. Godfrey, C., Muses, S., McClorey, G., Wells, K. E., Coursindel, T., Terry, R. L., Betts, C., Hammond, S., O’Donovan, L., Hildyard, J., El Andaloussi, S., Gait, M. J., Wood, M. J., and Wells, D. J. (2015) How much dystrophin is enough: the physiological consequences of different levels of dystrophin in the mdx mouse. Human molecular genetics 24, 4225–4237

28. Pearson, J., and Sabarra, A. (1974) A method for obtaining longitudinal cryostat sections of living muscle without contraction artifacts. Stain Technol 49, 143–146

29. Preibisch, S., Saalfeld, S., and Tomancak, P. (2009) Globally optimal stitching of tiled 3D microscopic image acquisitions. Bioinformatics 25, 1463–1465

30. Yen, J. C., Chang, F. J., and Chang, S. (1995) A new criterion for automatic multilevel thresholding. IEEE Trans Image Process 4, 370–378

31. Heintzmann, R., and Ficz, G. (2006) Breaking the resolution limit in light microscopy. Brief Funct Genomic Proteomic 5, 289–301

32. Milo, R., Jorgensen, P., Moran, U., Weber, G., and Springer, M. (2010) BioNumbers--the database of key numbers in molecular and cell biology. Nucleic acids research 38, D750–753

33. Hildyard, J. C. W., Finch, A. M., and Wells, D. J. (2019) Identification of qPCR reference genes suitable for normalizing gene expression in the mdx mouse model of Duchenne muscular dystrophy. PLoS One 14, e0211384

34. Hildyard, J. C. W., Taylor-Brown, F., Massey, C., Wells, D. J., and Piercy, R. J. (2018) Determination of qPCR Reference Genes Suitable for Normalizing Gene Expression in a Canine Model of Duchenne Muscular Dystrophy. J Neuromuscul Dis 5, 177–191

35. ACDbio. (2017) RNAscope Reference Guide.

36. Klein, C. J., Coovert, D. D., Bulman, D. E., Ray, P. N., Mendell, J. R., and Burghes, A. H. (1992) Somatic reversion/suppression in Duchenne muscular dystrophy (DMD): evidence supporting a frame-restoring mechanism in rare dystrophin-positive fibers. American journal of human genetics 50, 950–959

37. Sicinski, P., Geng, Y., Ryder-Cook, A. S., Barnard, E. A., Darlison, M. G., and Barnard, P. J. (1989) The molecular basis of muscular dystrophy in the mdx mouse: a point mutation. Science (New York,N.Y.) 244, 1578–1580

38. Chamberlain, J. S., Pearlman, J. A., Muzny, D. M., Gibbs, R. A., Ranier, J. E., Caskey, C. T., and Reeves, A. A. (1988) Expression of the murine Duchenne muscular dystrophy gene in muscle and brain. Science (New York, N.Y.) 239, 1416–1418

39. Maquat, L. E., Tarn, W. Y., and Isken, O. (2010) The pioneer round of translation: features and functions. Cell 142, 368–374

40. Bevilacqua, P. C., Ritchey, L. E., Su, Z., and Assmann, S. M. (2016) Genome-Wide Analysis of RNA Secondary Structure. Annu Rev Genet 50, 235–266

41. Lai, W. C., Kayedkhordeh, M., Cornell, E. V., Farah, E., Bellaousov, S., Rietmeijer, R., Salsi, E., Mathews, D. H., and Ermolenko, D. N. (2018) mRNAs and lncRNAs intrinsically form secondary structures with short end-to-end distances. Nat Commun 9, 4328

42. Yoffe, A. M., Prinsen, P., Gopal, A., Knobler, C. M., Gelbart, W. M., and Ben-Shaul, A. (2008) Predicting the sizes of large RNA molecules. Proceedings of the National Academy of Sciences of the United States of America 105, 16153–16158

43. Chen, H., Meisburger, S. P., Pabit, S. A., Sutton, J. L., Webb, W. W., and Pollack, L. (2012) Ionic strength-dependent persistence lengths of single-stranded RNA and DNA. Proceedings of the National Academy of Sciences of the United States of America 109, 799–804

44. Chi, Q. J., Wang, G. X., and Jiang, J. H. (2013) The persistence length and length per base of single-stranded DNA obtained from fluorescence correlation spectroscopy measurements using mean field theory. Physica A 392, 1072–1079

45. Milstein, J. N., and Meiners, J.-C. (2013) Worm-Like Chain (WLC) Model. In Encyclopedia of Biophysics (Roberts, G. C. K., ed) pp. 2757–2760, Springer Berlin Heidelberg, Berlin, Heidelberg

46. Petkova, M. V., Morales-Gonzales, S., Relizani, K., Gill, E., Seifert, F., Radke, J., Stenzel, W., Garcia, L., Amthor, H., and Schuelke, M. (2016) Characterization of a Dmd (EGFP) reporter mouse as a tool to investigate dystrophin expression. Skeletal muscle 6, 25

47. de Leon, M. B., Montanez, C., Gomez, P., Morales-Lazaro, S. L., Tapia-Ramirez, V., Valadez-Graham, V., Recillas-Targa, F., Yaffe, D., Nudel, U., and Cisneros, B. (2005) Dystrophin Dp71 expression is down-regulated during myogenesis: role of Sp1 and Sp3 on the Dp71 promoter activity. The Journal of biological chemistry 280, 5290–5299

48. Kawaguchi, T., Niba, E. T. E., Rani, A. Q. M., Onishi, Y., Koizumi, M., Awano, H., Matsumoto, M., Nagai, M., Yoshida, S., Sakakibara, S., Maeda, N., Sato, O., Nishio, H., and Matsuo, M. (2018) Detection of Dystrophin Dp71 in Human Skeletal Muscle Using an Automated Capillary Western Assay System. Int J Mol Sci 19, 1546

49. Siebrasse, J. P., Kaminski, T., and Kubitscheck, U. (2012) Nuclear export of single native mRNA molecules observed by light sheet fluorescence microscopy. Proceedings of the National Academy of Sciences of the United States of America 109, 9426–9431

50. Spencer, C. A., and Groudine, M. (1990) Transcription elongation and eukaryotic gene regulation. Oncogene 5, 777–785

51. Schoenberg, D. R., and Maquat, L. E. (2012) Regulation of cytoplasmic mRNA decay. Nature reviews. Genetics 13, 246–259

52. McCarthy, J. J., Andrews, J. L., McDearmon, E. L., Campbell, K. S., Barber, B. K., Miller, B. H., Walker, J. R., Hogenesch, J. B., Takahashi, J. S., and Esser, K. A. (2007) Identification of the circadian transcriptome in adult mouse skeletal muscle. Physiol Genomics 31, 86–95

53. Nicolas, D., Phillips, N. E., and Naef, F. (2017) What shapes eukaryotic transcriptional bursting? Mol Biosyst 13, 1280–1290

54. Newlands, S., Levitt, L. K., Robinson, C. S., Karpf, A. B., Hodgson, V. R., Wade, R. P., and Hardeman, E. C. (1998) Transcription occurs in pulses in muscle fibers. Genes Dev 12, 2748–2758

55. Cacchiarelli, D., Incitti, T., Martone, J., Cesana, M., Cazzella, V., Santini, T., Sthandier, O., and Bozzoni, I. (2011) miR-31 modulates dystrophin expression: new implications for Duchenne muscular dystrophy therapy. EMBO reports 12, 136–141

56. Cacchiarelli, D., Martone, J., Girardi, E., Cesana, M., Incitti, T., Morlando, M., Nicoletti, C., Santini, T., Sthandier, O., Barberi, L., Auricchio, A., Musaro, A., and Bozzoni, I. (2010) MicroRNAs involved in molecular circuitries relevant for the Duchenne muscular dystrophy pathogenesis are controlled by the dystrophin/nNOS pathway. Cell metabolism 12, 341–351

57. Fanin, M., Danieli, G. A., Vitiello, L., Senter, L., and Angelini, C. (1992) Prevalence of dystrophin-positive fibers in 85 Duchenne muscular dystrophy patients. Neuromuscular disorders: NMD 2, 41–45

58. Fanin, M., Danieli, G. A., Cadaldini, M., Miorin, M., Vitiello, L., and Angelini, C. (1995) Dystrophin-positive fibers in Duchenne dystrophy: origin and correlation to clinical course. Muscle & nerve 18, 1115–1120

59. Danko, I., Chapman, V., and Wolff, J. A. (1992) The frequency of revertants in mdx mouse genetic models for Duchenne muscular dystrophy. Pediatric research 32, 128–131

60. Lu, Q. L., Morris, G. E., Wilton, S. D., Ly, T., Artem’yeva, O. V., Strong, P., and Partridge, T. A. (2000) Massive idiosyncratic exon skipping corrects the nonsense mutation in dystrophic mouse muscle and produces functional revertant fibers by clonal expansion. The Journal of cell biology 148, 985–996

61. Pigozzo, S. R., Da Re, L., Romualdi, C., Mazzara, P. G., Galletta, E., Fletcher, S., Wilton, S. D., and Vitiello, L. (2013) Revertant Fibers in the mdx Murine Model of Duchenne Muscular Dystrophy: An Age-and Muscle-Related Reappraisal. PLOS ONE 8, e72147

62. Wilton, S. D., Dye, D. E., and Laing, N. G. (1997) Dystrophin gene transcripts skipping the mdx mutation. Muscle & nerve 20, 728–734

63. Verhaart, I. E. C., van Vliet-van den Dool, L., de Sipkens, J. A., Kimpe, S. J., Kolfschoten, I. G. M., van Deutekom, J. C. T., Liefaard, L., Ridings, J. E., Hood, S. R., and Aartsma-Rus, A. (2014) The Dynamics of Compound, Transcript, and Protein Effects After Treatment With 2OMePS Antisense Oligonucleotides in mdx Mice. Molecular therapy Nucleic acids 3, e148–e148

64. Amoasii, L., Hildyard, J. C. W., Li, H., Sanchez-Ortiz, E., Mireault, A., Caballero, D., Harron, R., Stathopoulou, T.-R., Massey, C., Shelton, J. M., Bassel-Duby, R., Piercy, R. J., and Olson, E. N. (2018) Gene editing restores dystrophin expression in a canine model of Duchenne muscular dystrophy. Science (New York, N.Y.) 362, 86–91

65. Long, C., Amoasii, L., Mireault, A. A., McAnally, J. R., Li, H., Sanchez-Ortiz, E., Bhattacharyya, S., Shelton, J. M., Bassel-Duby, R., and Olson, E. N. (2016) Postnatal genome editing partially restores dystrophin expression in a mouse model of muscular dystrophy. Science (New York, N. Y.) 351, 400–403

66. Amoasii, L., Long, C., Li, H., Mireault, A. A., Shelton, J. M., Sanchez-Ortiz, E., McAnally, J. R., Bhattacharyya, S., Schmidt, F., Grimm, D., Hauschka, S. D., Bassel-Duby, R., and Olson, E. N. (2017) Single-cut genome editing restores dystrophin expression in a new mouse model of muscular dystrophy. Sci Transl Med 9

67. Nelson, C. E., Hakim, C. H., Ousterout, D. G., Thakore, P. I., Moreb, E. A., Castellanos Rivera, R. M., Madhavan, S., Pan, X., Ran, F. A., Yan, W. X., Asokan, A., Zhang, F., Duan, D., and Gersbach, C. A. (2016) In vivo genome editing improves muscle function in a mouse model of Duchenne muscular dystrophy. Science (New York, N. Y.) 351, 403–407

68. Tabebordbar, M., Zhu, K., Cheng, J. K. W., Chew, W. L., Widrick, J. J., Yan, W. X., Maesner, C., Wu, E. Y., Xiao, R., Ran, F. A., Cong, L., Zhang, F., Vandenberghe, L. H., Church, G. M., and Wagers, A. J. (2016) In vivo gene editing in dystrophic mouse muscle and muscle stem cells. Science (New York, N.Y.) 351, 407–411

