## Supplementary figures and images for "Multiplex *in situ* hybridization within a single transcript: RNAscope reveals dystrophin mRNA dynamics"

### Supplementary Figure 1

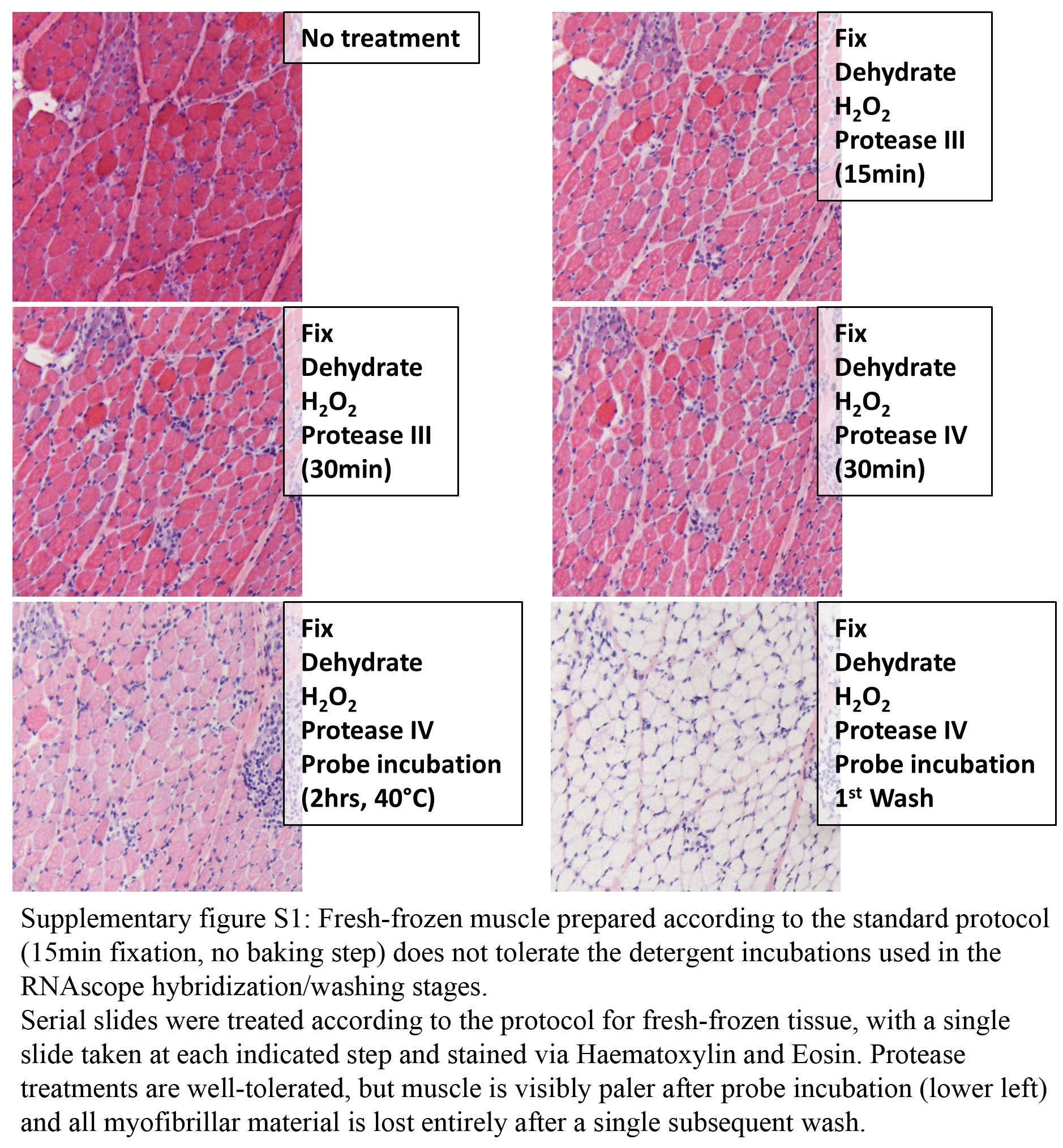

### Supplementary Figure 2

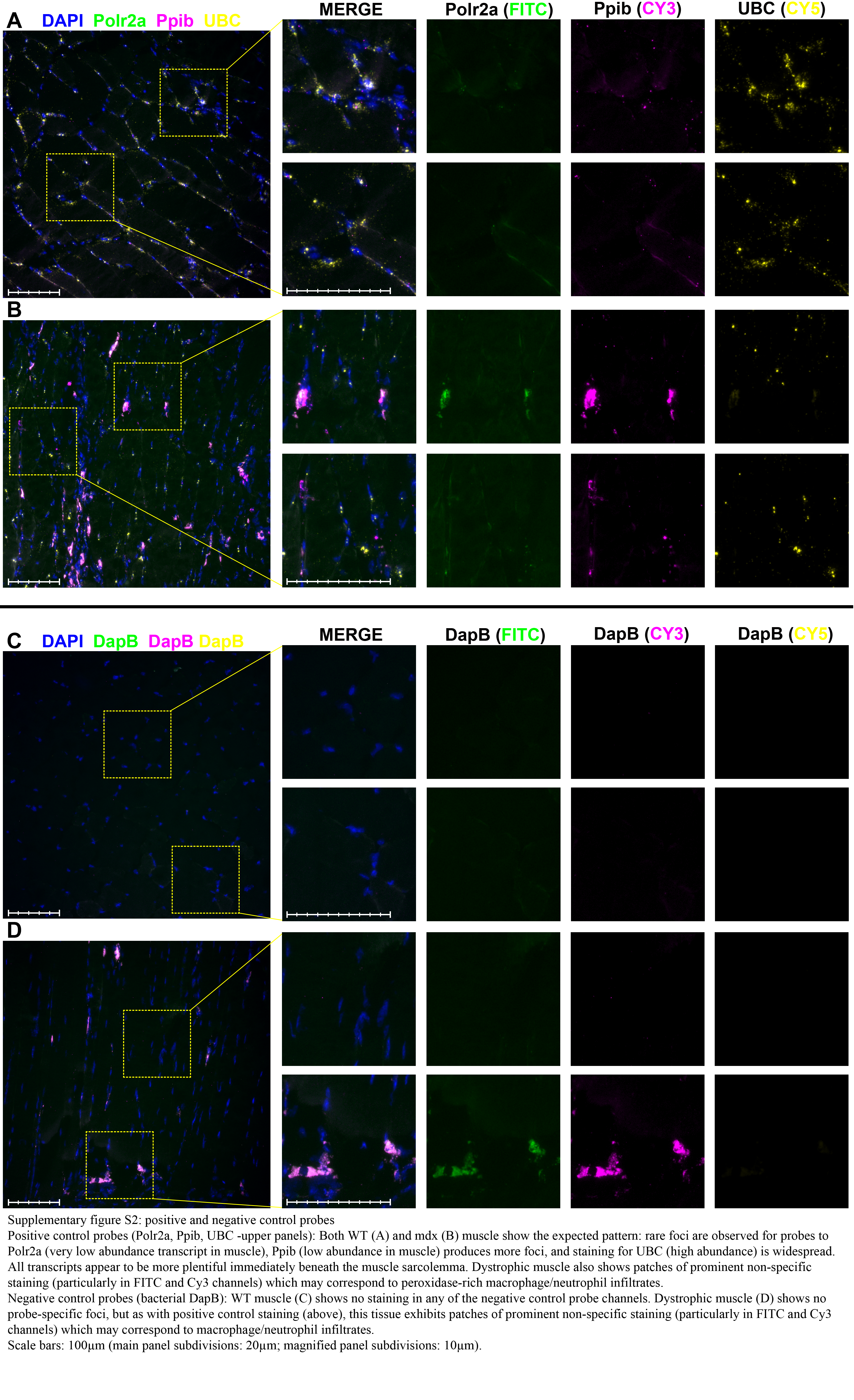

### Supplementary Figure 3

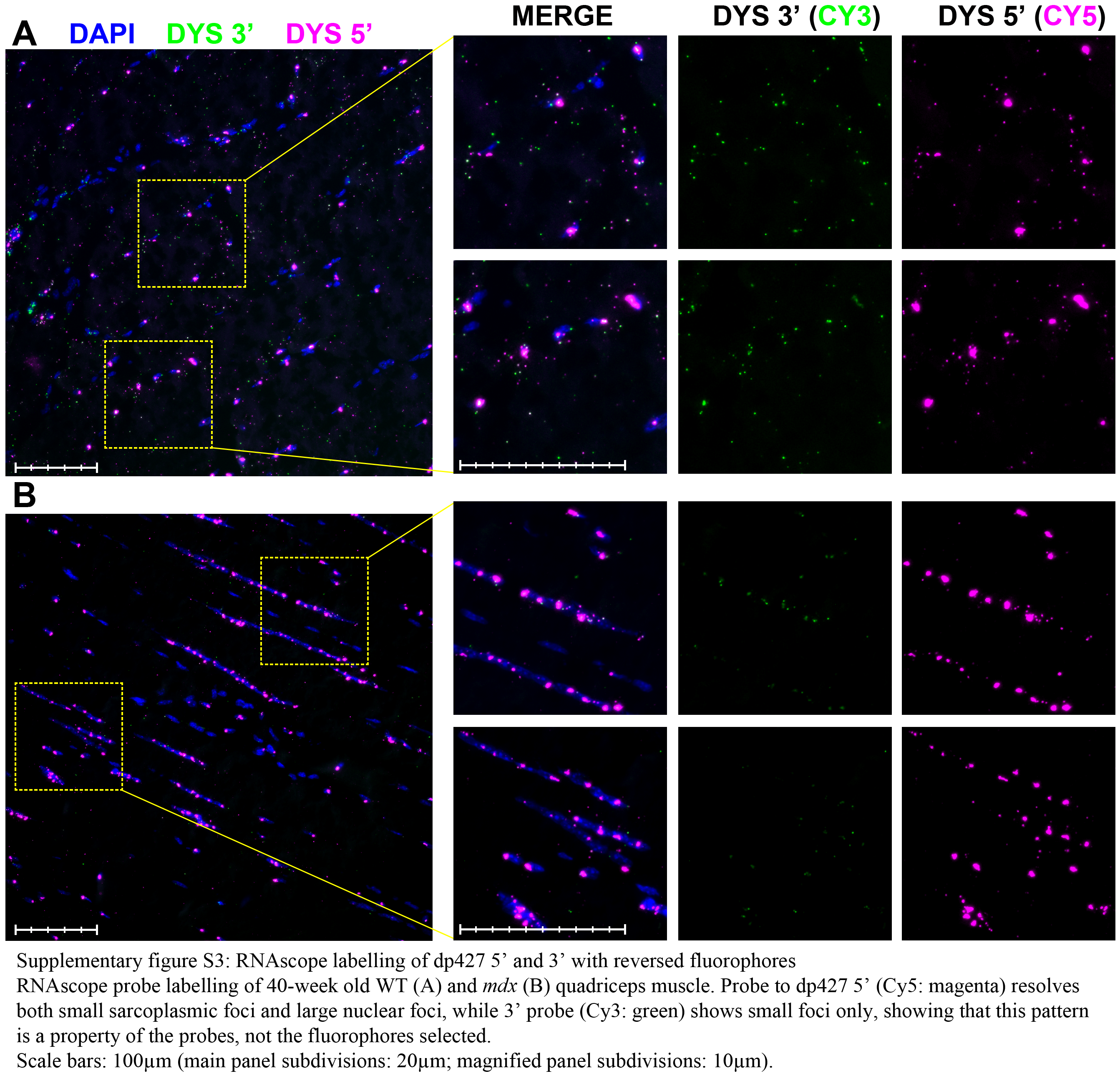

### Supplementary Figure 4

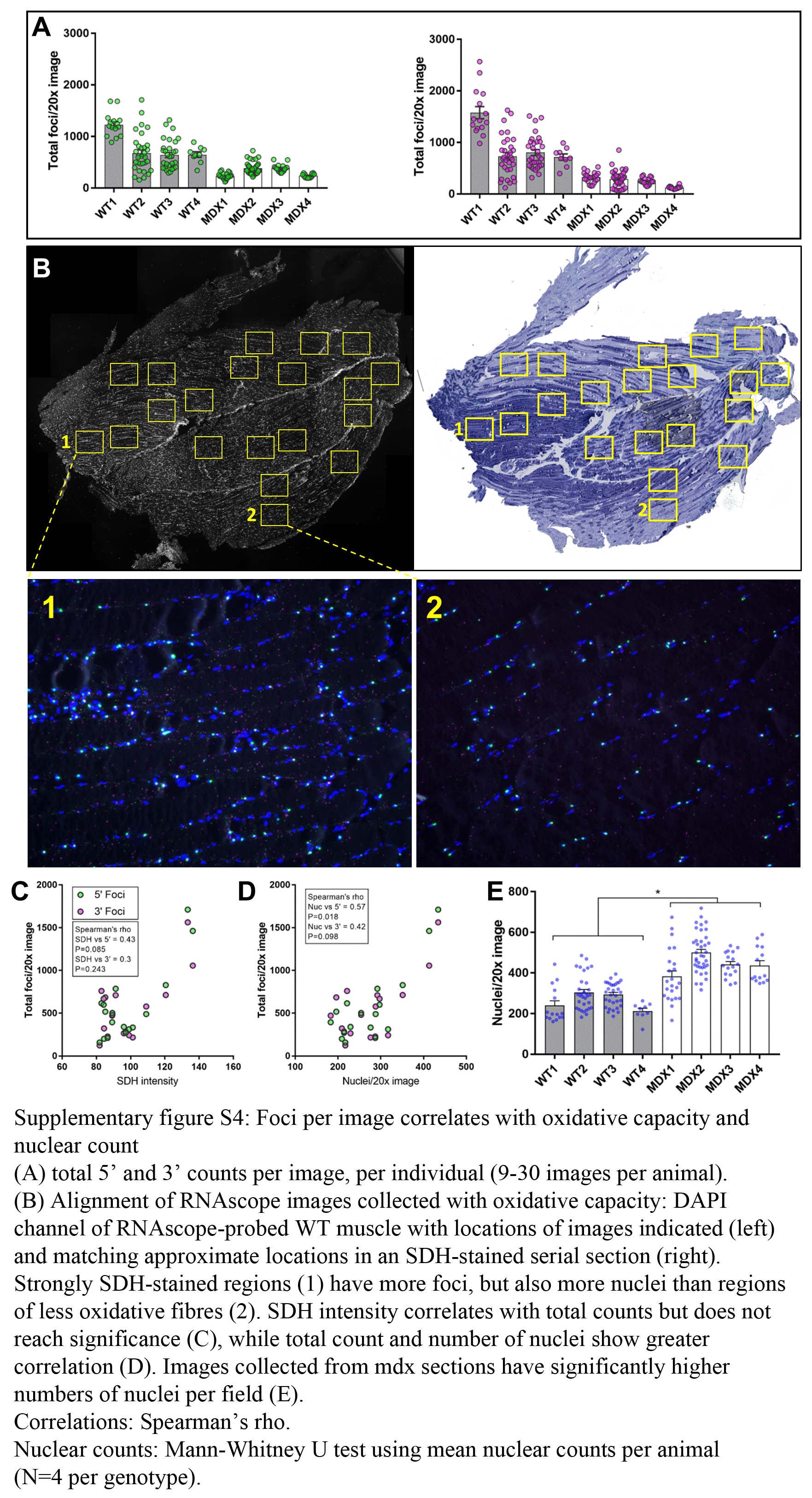

### Supplementary Figure 5

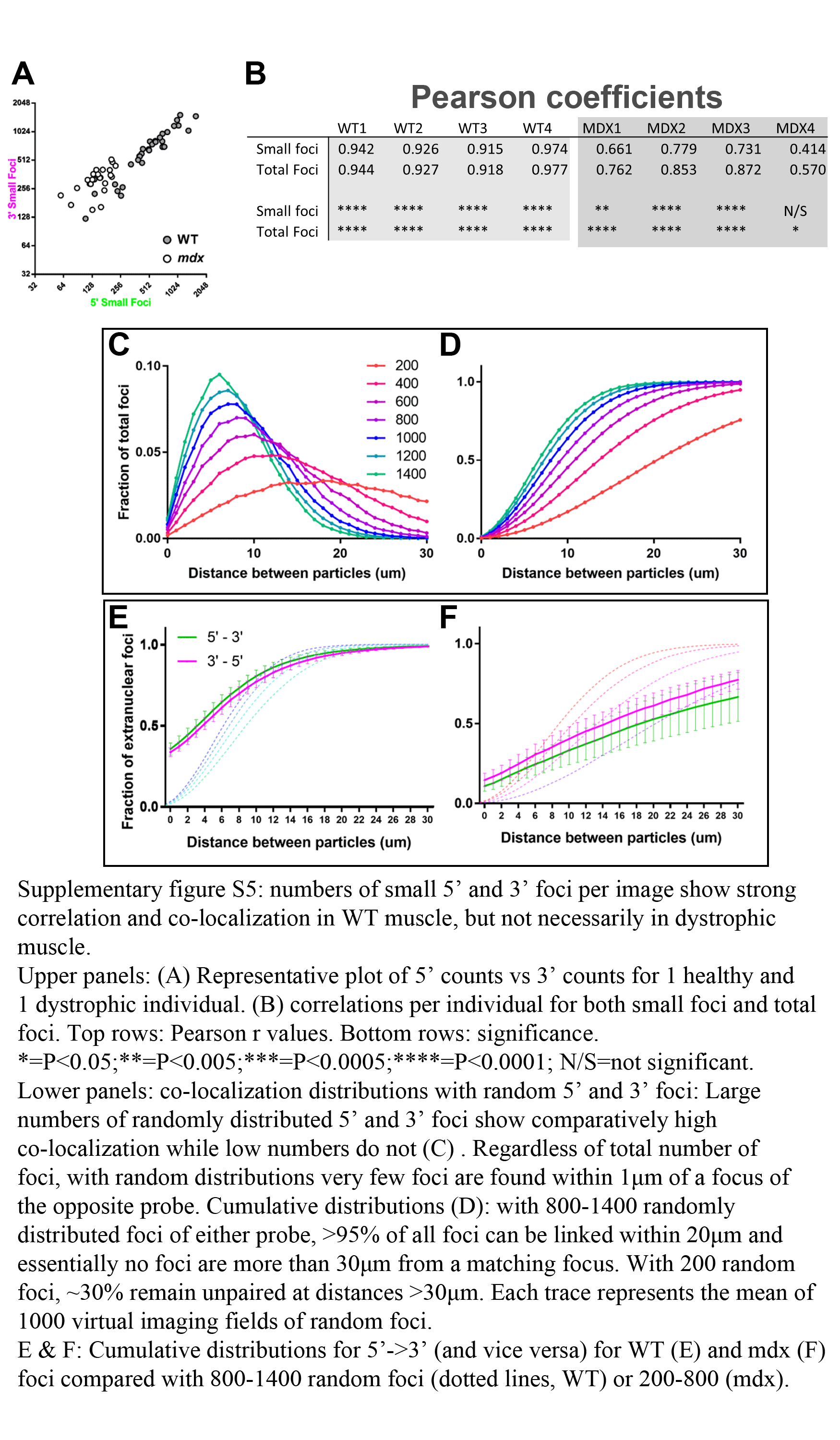

### Supplementary Figure 6

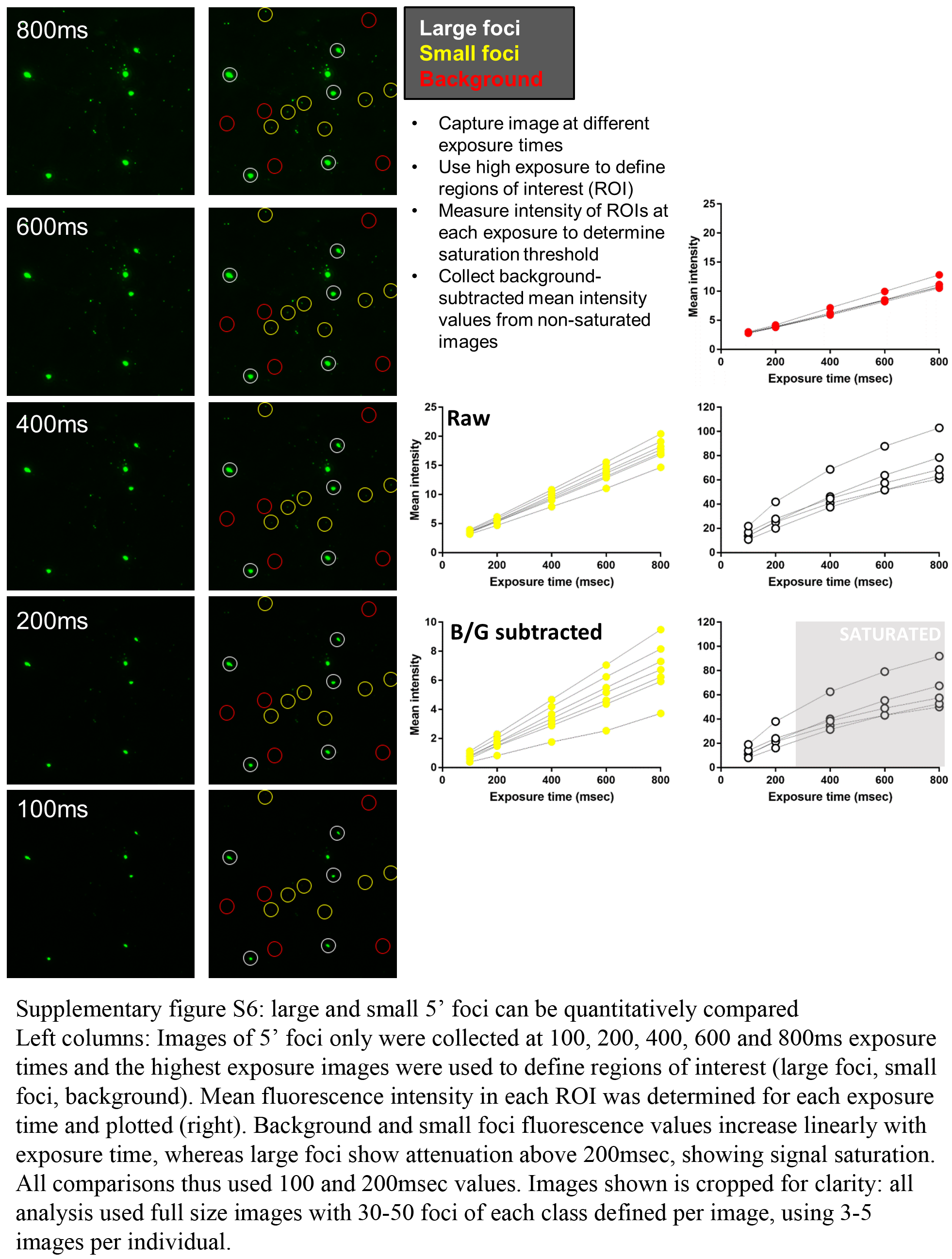

### Supplementary Figure 7

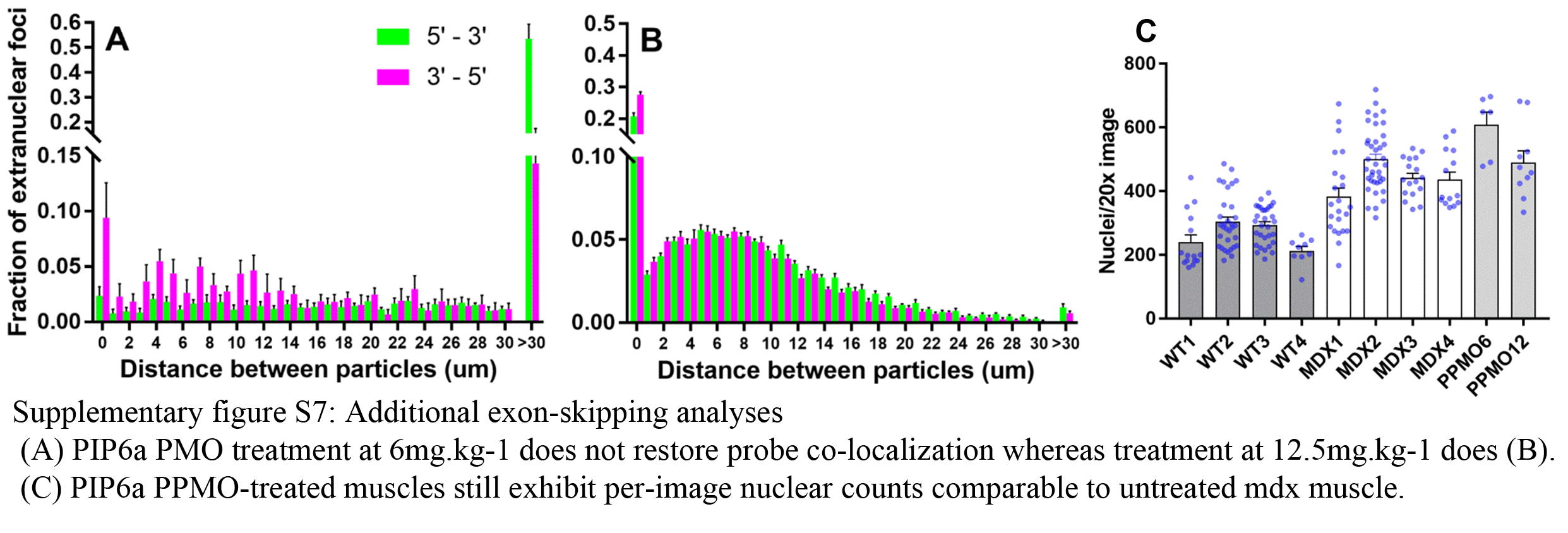

### Supplementary Figure 8

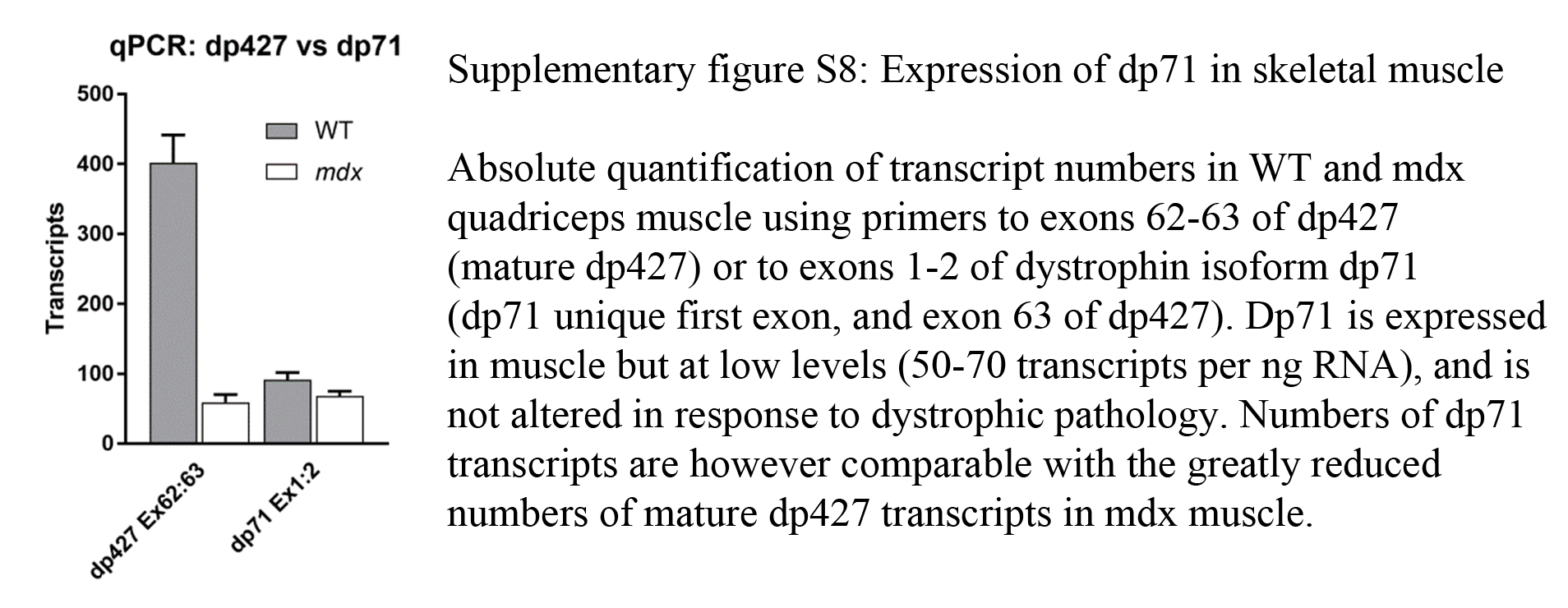
